# A neural circuit basis for binasal input-enhanced chemosensory avoidance

**DOI:** 10.1101/2021.02.20.431946

**Authors:** Samuel K. H. Sy, Danny C. W. Chan, Roy C. H. Chan, Jing Lyu, Zhongqi Li, Kenneth K. Y. Wong, Chung Hang Jonathan Choi, Vincent C. T. Mok, Hei-Ming Lai, Yu Hu, Ho Ko

## Abstract

Our understanding of how bilaterian animals utilize parallel input channels from paired sensory organs to optimize chemosensory behavior and the underlying neural circuit mechanisms are limited. Here we developed microfluidics-based behavioral and brainwide imaging platforms to study the neural integration of binasal inputs and chemosensory avoidance in larval zebrafish. We show that larval zebrafish efficiently escape from cadaverine-carrying streams by making more frequent swim bouts and larger undirected turns. Binasal inputs are strictly required for the nasal input-dependent component of klinokinesis, while each nasal input additively enhances angular orthokinesis. Throughout brain regions, including those along the olfactory processing pathways, a distributed neural representation with a wide spectrum of ipsilateral-contralateral nasal stimulus selectivity is maintained. Nonlinear sensory information gain with bilateral signal convergence is especially prominent in neurons weakly encoding unilateral cadaverine stimulus, and associated with stronger activation of sensorimotor neurons in the downstream brain regions. Collectively, these results provide insights into how the vertebrate model sums parallel input signals to guide chemosensory avoidance behavior.

## Introduction

Bilaterian animals, which constitute the vast majority of animal species, have paired and symmetrical sensory organs. Integration of paired sensory organ inputs across different sensory modalities have been previously studied in humans^1–3^, mammals^4–8^ and insects^9,10^. In principle, such a design offers several potential advantages. For example, inputs from spatially segregated, parallel sensory channels may be summed (or averaged) to enhance the signal-to-noise ratio of cue detection^11^, or used in stereo comparison for spatial information extraction^12,13^.

For chemosensory behaviors, the role of bilateral sensory input appears to be species- and context-dependent. Humans and rodents can employ a strategy combining sequential sniffing and stereo comparison to efficiently localize odorant sources^1,14–17^. In *Drosophila* larvae, bilateral dorsal olfactory organ inputs improve odor gradient detection during serial sampling to guide navigation^11^. Adult *Drosophila* instead appear capable of stereo comparison between olfactory antenna inputs for odorant localization^18^. Similarly, a dependence of chemotaxis on bilateral olfactory inputs has been reported in some insects, such as honey bees and ants^9,10^. For aquatic animals, sampling waterborne chemical cues is essential for survival. Despite well-known chemosensory avoidance and attraction behaviors in various aquatic species^19–24^, how binasal inputs are utilized in guiding these behaviors has been scarcely reported. To date, the only behavioral report pertains to *Mustelus canis*, which can compare inter-nasal odor arrival times to bias turns towards a food cue^25^.

Moreover, the neural basis of processing parallel inputs from paired sensory organs for the instruction of motor output is incompletely understood. In particular, brainwide level bilateral sensory integration and sensorimotor transformation underlying chemosensory behavior remains virtually unknown. While interhemispheric projections arising from early sensory pathways or higher-order areas are well-described in many species^26,27^, detailed circuit mechanisms of how these pathways mediate sensory processing are only beginning to be unraveled^28–32^. Limited by tools available for studying the relatively large brains of numerous species, past works mostly focused on individual regions, such as the binocular visual areas^32–35^, the primary auditory cortex^12,36–40^ or restricted areas of the olfactory processing pathways^41,42^. However, both the integration of bilateral sensory input and subsequent sensorimotor transformation may occur on a distributed, brainwide scale. Near-complete sampling of whole-brain neuronal dynamics during sensory processing and behavior is therefore a powerful approach to gain more in-depth understandings of the underlying circuit mechanisms^43–45^.

The zebrafish olfactory system shares a conserved general circuit layout as other vertebrates, and offers the distinct advantage of permitting optical imaging of relatively large proportions of neurons along the early olfactory pathways^46^. It is thus an attractive model system for studying the computational principles of sensory processing^46^. Remarkably, within one-week post-fertilization, larval zebrafish already have a rich repertoire of innate behaviors. These include chemosensory behaviors, alongside phototaxis, oculomotor and optokinetic responses, prey capture and various escape responses^45,47^. With a relatively small, transparent brain that comprises a more tractable number of neurons (~10^5^), much has been learnt about the circuitries mediating sensory processing and sensorimotor behaviors in larval zebrafish^29,48–66^. Larval zebrafish also offers a unique opportunity for studying the brainwide organizational principles of neural circuits mediating bilateral sensory input integration. For example, circuit processing principles for the binocular visual input-driven optomotor response^66^ and prey capture^29^ behaviors have been investigated. For the olfactory system, interhemispheric crosstalks between parallel olfactory channels have been shown at the anatomical level^67,68^. However, functional data are needed to constrain circuit models with respect to properties such as projection pathway dominance, integration sites, and the rules of input summation by individual neurons.

In this work, we use larval zebrafish as the model organism to study the neural basis of parallel nasal input integration in chemosensory avoidance. We first develop a laminar flow-based assay with a static chemical border, to investigate the avoidance behavior of larval zebrafish upon naturalistic encounters of an aversive stimulus. Using an odorant (cadaverine) which does not elicit strong escape responses yet is survival-relevant to avoid, we demonstrate behavioral changes with significant consequences on escape efficiency that differentially depend on the integrity of bilateral olfactory placodes (OPs). We also develop an integrated microfluidics-light sheet imaging system to interrogate whole-brain neuronal dynamics during spatiotemporally precise unilateral or bilateral OP stimulation in partially restrained larval subjects, which could not be achieved by previously reported experimental platforms^69–73^. This approach enables us to reveal a distributed neural representation of cadaverine sensing, characterized by a wide range of ipsilateral-contralateral <u>nasal</u> input selectivity throughout brain regions along the rostral-caudal axis, and the convergence of bilateral afferent signals on a brainwide scale. We show that sensory information gain with bilateral signal convergence is generally supralinear, and such synergistic input summation is associated with stronger activation of sensorimotor and motor output-encoding neuronal subsets.

## Results

### Larval zebrafish requires binasal inputs to efficiently escape cadaverine zone

In the natural environment, a fish needs to navigate away from noxious chemicals. To study chemosensory avoidance under a mimicking condition, we developed a microfluidics-based setup that maintains a precisely defined spatio-chemical landscape. The setup employs a rectangular swimming arena with three fluid inlets and two fluid outlets (**Figure 1A**). Computational fluid dynamics simulation shows laminar flow at steady state with very low flow velocity profile over the majority of the area except near inlets and outlets (**Supplementary Figure 1A**). The laminar flow enables the maintenance of static boundaries (**Supplementary Figure 1B**) with steep chemical concentration gradients (**Supplementary Figure 1C**) between fluidic stream zones (**Figure 1A**). Upon crossing by a larva, the border is transiently disturbed yet quickly restored within a few seconds (**Supplementary Figure 1D**). During an avoidance assay, three to six zebrafish larvae aged 5–7 days post-fertilization (d.p.f.) freely swim in the arena with a noxious chemical zone in the absence of visible light (**Figure 1A**). We carried out high-speed imaging (240 Hz frame rate) under infrared (IR) illumination and tracking of larval zebrafish head orientation and coordinates for the quantification of avoidance behavior (**Figure 1A**, see **Methods**). With a static chemical zone boundary, this assay allowed us to repeatedly sample events of entering and escaping the well-defined noxious chemical zone. Assays with water streams in all zones served as control.

**Figure 1.**
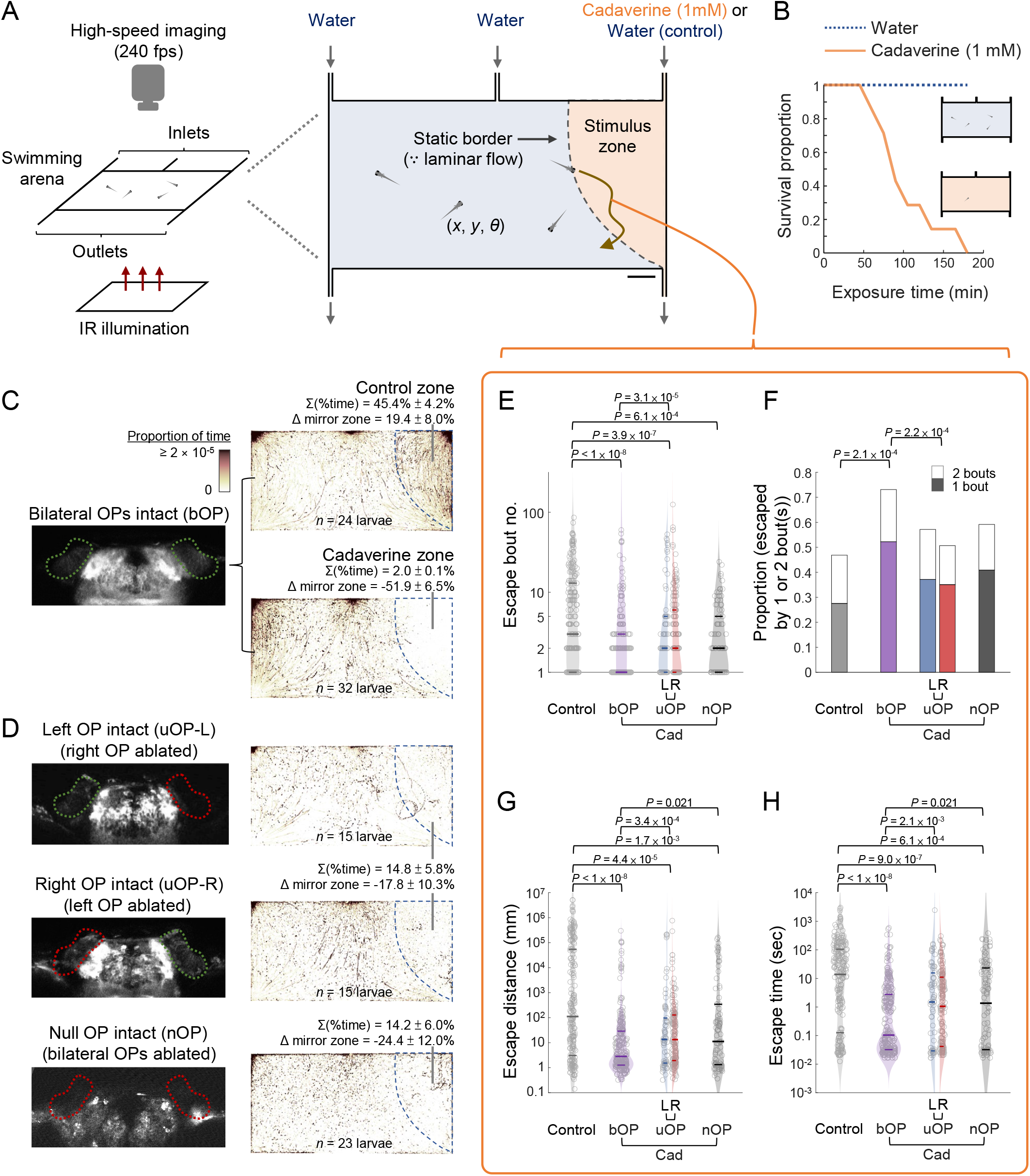
Larval zebrafish performance in a fluidics-based avoidance behavioral assay. **(A)** Schematics of the avoidance behavioral assay. Zebrafish larvae swimming in a two-dimensional arena (60 mm × 30 mm × 1.5 mm) are imaged at high speed (240 fps) under infrared (IR) illumination in the absence of visible light. A noxious chemical zone (stimulus zone) is created and maintained by a constant slow inflow of cadaverine (1 mM in water) via the rightmost fluid inlet. Assays in which all zones are filled by water streams serve as control. The laminar flow maintains a static border between the zones. The coordinates (*x*, *y*) and orientation (*θ*) of the center of the head are tracked and analysed for each larval zebrafish. Scale bar: 0.5 cm. **(B)** Proportion of larvae surviving in an arena shown in **(A)** filled by just cadaverine (1 mM in water, *n* = 7, orange solid line) or just water (control, *n* = 6, blue dotted line) with time. **(C)** Left panel: Two-photon image of a zebrafish larva with the olfactory placodes (OPs) outlined by green dotted lines (designated as bilateral OP-intact, or bOP larvae). Right panels: Footprints of bOP larvae, expressed in proportion of time occupied by head center coordinates for each pixel, over 2-hour behavioral assays with all zones filled by water streams (upper panel, with the rightmost zone outlined by dashed line), or the stimulus zone filled by cadaverine stream (lower panel, with the cadaverine zone outlined by dashed line). **(D)** Left panels: Images of example unilateral OP-intact (uOP) larvae with ablation of the right OP (upper panel, left OP intact or uOP-L) or left OP (middle panel, right OP intact or uOP-R), and a larva with ablation of both OPs (lower panel, null OP intact or nOP). Intact OP(s) are outlined in green, and ablated OPs are outlined in red dotted lines. Right panels: Footprints of the different larvae groups over 2-hour avoidance assays (cadaverine zone outlined by dashed line). In **(C)** and **(D)**, the percentages of time spent in the stimulus zone, their differences from that in the mirror water zone (Δ mirror zone, a negative value indicates less time spent in the stimulus zone) are shown (with SEMs across trials). **(E)** Bout number, **(F)** proportion of entry-to-exit events with only 1 or 2 bouts (*P*-value: Chi-squared test with Tukey’s post-hoc test, comparing 2-bout event proportions), **(G)** distance travelled, and **(H)** time taken to escape the stimulus zone in control assays (bOP larvae in water-only arena) and avoidance behavioral assays (bOP larvae, left OP-intact (L) or right OP-intact (R) uOP larvae, or nOP larvae in arenas with cadaverine stream in the stimulus zone). In **(E)**, **(G)** and **(H),** the parameters are plotted in log scales. Horizontal lines indicate the medians, 75 and 25 percentiles for each group. *P*-values: Kruskal–Wallis test with Tukey’s post-hoc test.

Cadaverine is a diamine product of putrefaction of corpses. Apart from being an alarm cue, we observed that it is toxic to larval zebrafish, as prolonged exposure to cadaverine (1 mM) in a survival assay invariably led to death (**Figure 1B**, see **Methods**). However, simply inspecting individual events of encountering cadaverine in the avoidance assay, the larvae appeared incapable of performing immediate, strong escape, in line with a previous report^74^. Yet, during more prolonged assay (2 hours), the zebrafish larvae exhibited clear avoidance of the cadaverine zone, as they spent substantially less time in the cadaverine zone than the spatially identical stimulus zone filled by water stream (control zone) (**Figure 1C**). This indicated that cadaverine avoidance in the larval zebrafish arises from more subtle behavioral changes that require sufficient observations to reveal. To test the dependence of the avoidance behavior on the integrity of the OPs, we performed two-photon laser ablation of unilateral or bilateral OPs and carried out the assay in three additional groups of larvae (i.e., either the left, the right, or both OPs were ablated, **Figure 1D**). Interestingly, although the unilateral (uOP) and null OP-intact (nOP) larvae could still avoid the cadaverine zone, the proportions of time spent in the cadaverine zone were higher than the bilateral OP-intact (bOP) larvae (**Figures 1C and 1D**).

To compare escape efficiency from the noxious odor zone, we examined several avoidance performance metrics, including number of swim bouts (as zebrafish larvae swim in discrete bouts), time and distance travelled to escape, for each cadaverine zone entry event (**Figures 1E–1H**). The bOP larvae escaped from the cadaverine zone with fewer bouts than the uOP larvae (**Figure 1E**), and could escape using only one or two bouts in over 70% of the events, while the corresponding proportions for the uOP larvae in cadaverine zone were only slightly higher than that of the bOP larvae in control zone (**Figure 1F**). Consequently, the bOP larvae on average travelled significantly shorter distances (**Figure 1G**) and took less time (**Figure 1H**) to leave the cadaverine zone than the uOP larvae. Consistent with the observation that uOP and nOP larvae spent similar proportions of time in the cadaverine zone (**Figure 1D**), these parameters did not differ between the uOP and nOP larvae groups (**Figures 1E–1H**). In larval zebrafish, the performance of chemosensory avoidance therefore depends on the integrity of binasal inputs, as either unilateral input alone did not enhance avoidance beyond that mediated by non-OP-dependent cadaverine detection. In the mirror zone of the arena (filled by water), all the OP-ablated groups (i.e., uOP and nOP) exhibited no significant differences in these parameters from the intact (i.e., bOP) larvae groups (**Supplementary Figures 2A–2D**). These confirmed the behavioral modulation by cadaverine, and showed that spontaneous swimming behaviors were not affected by the ablation procedure.

### Binasal inputs modulate turning behavior in cadaverine avoidance

In theory, different swim bout or turn kinematic changes could account for the observed bilateral OP-dependent efficiency of cadaverine avoidance. These include increase in swim bout frequency (i.e., klinokinesis), and angular or linear velocity (i.e., angular or linear orthokinesis) (**Figure 2A**). To understand the strategy employed by larval zebrafish, we analysed swim bout kinematic parameters for the different groups in control and cadaverine zones (**Figures 2B–2D**). During the assay, occasionally the larvae came into contact with each other or the arena boundary and elicited touch-initiated responses, or performed mechanosensory-based rheotaxis near the fluid inlets and outlets where fluid velocities were higher (**Supplementary Figure 1A**). As we focused on chemical-induced behaviors, the trajectories of these events were excluded from analysis on swim bout kinematics (see **Methods**).

**Figure 2.**
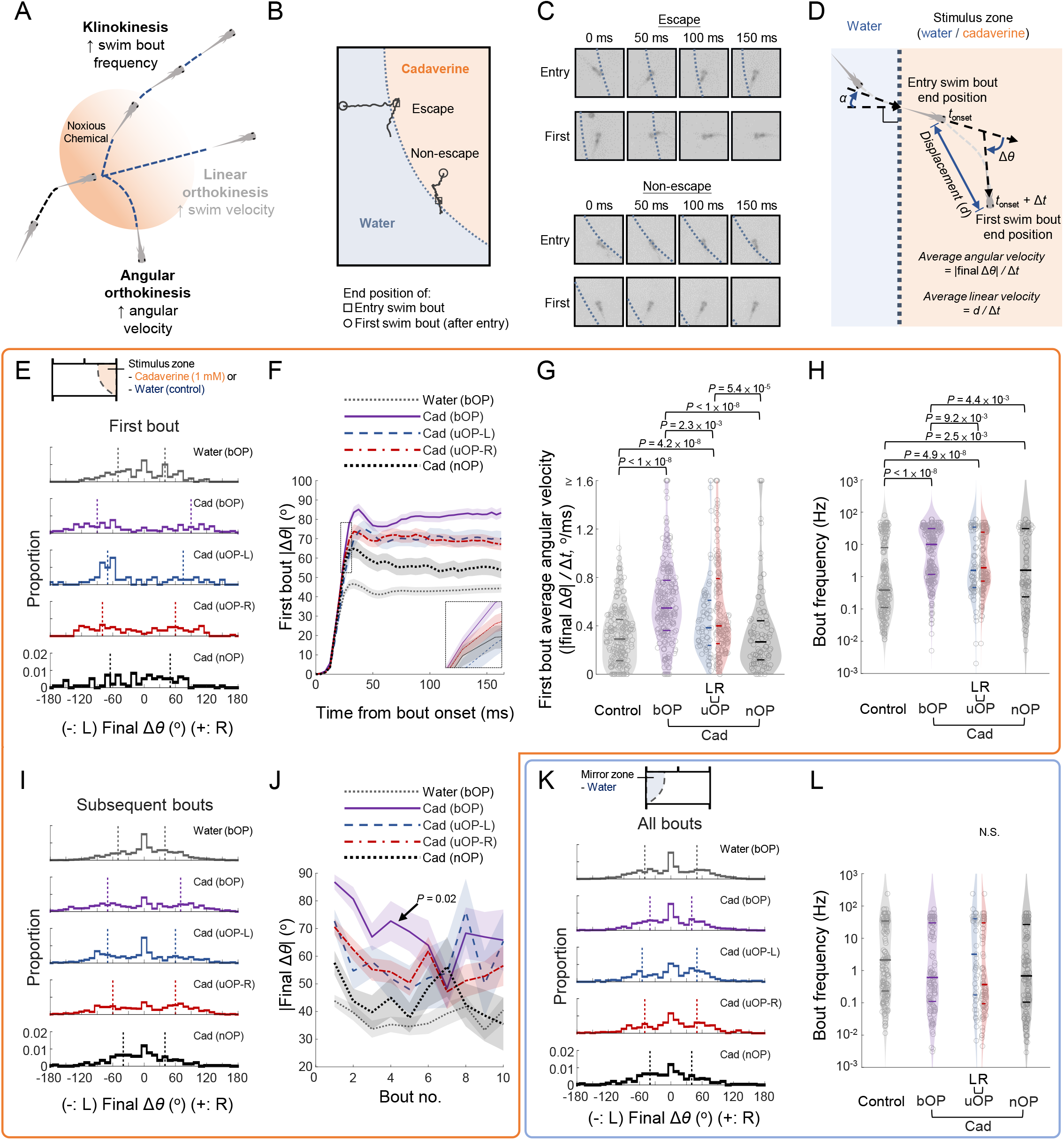
Kinematic parameters of swim bouts after entering stimulus zone. **(A)** Illustration of navigational strategies that may be adopted after encountering a noxious chemical, including klinokinesis, angular orthokinesis and linear orthokinesis. **(B)** Two example entry trajectories by a larval zebrafish, whereby the first swim bouts after entering the cadaverine zone resulted in escape (upper example) or further navigation into the region (lower example). **(C)** Snapshots of the tracked larval zebrafish at different time points after onset of the corresponding swim bouts (entry or first bout) in **(B)**. Dotted lines: cadaverine zone border. **(D)** Schematics of kinematic parameters extracted from first bouts, including incident angle (relative to vector normal to border) on stimulus zone entry (*α*), first bout onset time after stimulus zone entry (*t*_onset_), time-dependent change in head orientation (Δ*θ*), bout duration (Δ*t*), final orientation change on bout completion (final Δ*θ*), displacement, average angular and linear velocities. | | denotes absolute value. **(E) – (G)** Kinematic parameters of the first bouts after cadaverine encounter. **(E)** Histograms of turn angle distributions of first bouts (i.e., final Δ*θ*). **(F)** Dynamics of |Δ*θ*| after first bout onset (line: mean; shadow: SEM). Inset in **(F)** corresponds to the region outlined by the dashed box. **(G)** First bout average angular velocity (i.e., final Δ*θ*| / Δ*t*). **(H)** Swim bout frequency quantified from all stimulus zone entry and escape trajectories (plotted in log scale). **(I)** Histograms of turn angle distributions of subsequent bouts in the stimulus zone. **(J)** |Final Δ*θ*| vs. bout number (line: mean; shadow: SEM) in the stimulus zone. *P*-value: Mann-Kendall trend test. **(K)** Histograms of turn angle distributions of all bouts in the mirror zone. **(L)** Swim bout frequency in the mirror zone (plotted in log scale). In all plots, data are shown for control assays (bilateral OP-intact (bOP) larvae in water-only arena) and avoidance behavioral assays (bOP larvae, left (L) or right (R) unilateral OP-intact (uOP), or null OP-intact (nOP) larvae in arena with cadaverine (Cad) in the stimulus zone). In **(E), (I) and (K)**, dotted lines indicate the median angles in the corresponding directions. In **(G), (H) and (L)**, horizontal lines indicate the medians, 75 and 25 percentiles for each group. *P*-values: Kruskal–Wallis test with Tukey’s post-hoc test. N.S.: Non-significant.

Prompted by the observation that the bOP larvae could escape within two swim bouts more frequently than the uOP larvae (**Figure 1F**), we first focused on the initial swim bouts after encountering cadaverine (**Figures 2B–2D**). Following border crossing (i.e., stimulus zone entry), there were no differences in first swim bout onset times across the groups (**Supplementary Figure 2E**). However, the bOP larvae made larger turns during the first swim bouts in the cadaverine zone compared to the control zone, with a median angular magnitude difference approximately twice as large as that of the uOP larvae in cadaverine zone (**Figures 2E and 2F**). The angular magnitude difference was due to a higher angular velocity of the first bouts by the bOP larvae (**Figure 2G**), but not longer bout durations (**Supplementary Figure 2F**). The uOP larvae made larger and faster first turns than the nOP larvae in the cadaverine zone (**Figures 2E–G**), while the nOP larvae only made slightly larger first turns in the cadaverine zone (**Figures 2E and 2F**) with no statistically significant changes in angular velocity compared to the bOP group in control zone (**Figure 2G**). Nasal inputs therefore modulate the initial swim bouts in a graded manner, whereby each nasal input additively enhances angular orthokinesis. Note the highly overlapping distributions of the kinematic parameters, indicating behavioral changes that manifested only statistically that nevertheless significantly impacted avoidance performance.

We then analysed both initial and subsequent swim bouts during the escape journeys. We noted that all larvae groups exhibited higher swim bout frequency in the cadaverine zone compared to the bOP larvae in the control zone (**Figure 2H**). The nasal input-dependent component of the klinokinetic behavior strictly requires binasal inputs to manifest, as the bOP larvae increased their swim bout frequency the most, while there was no difference between the uOP and nOP groups (**Figure 2H**). These also suggested that klinokinesis is partially mediated by non-olfactory (e.g., gustatory) cadaverine chemosensation. Interestingly, we observed binasal input-dependent adaptation of swim bout angular magnitude upon continuous cadaverine exposure. Although both bOP and uOP larvae maintained higher angular magnitudes in the cadaverine zone than bOP larvae in the control zone on subsequent swim bouts (**Figure 2I**), the angular magnitude of bouts bOP larvae made converged onto that of uOP larvae with increasing bout number in the cadaverine zone (**Figure 2J**).

We did not find the larvae performing significant linear orthokinesis or directional turns to evade cadaverine. Only minimal differences in swim bout linear velocity in the cadaverine zone was observed between the groups (**Supplementary Figures 2G and 2H**). If larval zebrafish were capable of stereo comparison and lateralization of odor, the bOP larvae should be able to make directional turns depending on odor zone border-crossing angle, while the uOP larvae should bias their turns towards the ablated side upon encountering noxious odor. However, neither phenomenon was present (**Supplementary Figures 2I and 2J**). Importantly, we verified that swim bout angular changes (**Figure 2K**) and frequency (**Figure 2L**) exhibited no differences among the larvae groups in the mirror water zone. The decrease in angular magnitudes of consecutive swim bouts after leaving the cadaverine zone were also similar for all larvae groups (**Supplementary Figure 2K**). The ablation procedures therefore did not alter baseline swim bout kinematics.

Collectively, we concluded that under the assay, larval zebrafish perform faster and larger undirected turns more frequently, to efficiently escape the noxious cadaverine zone. Although unilateral nasal input and non-olfactory cadaverine detection can still modulate swimming behavior, binasal inputs are indispensable to optimize avoidance behavior with klinokinesis and large-magnitude angular orthokinesis that exhibits adaptation.

### Whole-brain neuronal imaging with spatiotemporally precise olfactory placode stimulation

Studying binasal input integration in larval zebrafish requires sequential presentations of unilateral and bilateral OP stimuli, which poses a unique methodological challenge due to the spatial proximity of the OPs (~100 μm apart). Since this could not be accomplished by previously reported techniques^69–73^, we custom-developed a poly(dimethylsiloxane) (PDMS)-based microfluidic device for partially restraining a larval zebrafish and directed chemical delivery to one or both OPs, that is fully compatible with simultaneous whole-brain imaging by light sheet fluorescence microscopy^75–77^ (**Figure 3**). The major functional compartments of the device include sets of chemical delivery microchannels, a head and waist-trapping chamber with a vertical and smooth side wall to ensure undisrupted propagation of incident excitation laser sheet (**Figure 3A and Supplementary Figure 3A,** also see **Methods**), and a tail chamber under IR illumination for the recording of tail flicking behavior at 200 Hz (**Figure 3B**). To prevent unwanted chemical spillover (**Supplementary Figure 3B**), we adopted a design whereby valve-gated fluid streams delivered from each side contain a central chemical stream which is insulated from the OP by the posterior water stream (**Figure 3C, Supplementary Figures 3B and 3C, *left***). Spatiotemporally precise chemical stimulus is delivered by turning off the insulating water stream(s), thereby allowing the chemical stream(s) to contact the respective OP(s), and terminated by turning off the chemical stream(s) (**Figure 3C, Supplementary Figure 3C, *middle* and *right***). Stimulus specificity quantified by the ratio of peak fluorescence intensity change from baseline (*dF/F*) within the nasal cavities using sodium fluorescein (10 and 100 μM) was invariably above 20:1 (intended:unintended side) across trials (**Figure 3C, *right***). In functional imaging experiments, the stimulus profiles visualized by a low concentration of sodium fluorescein (1 μM) were characterized by a rapid rise reaching a plateau (0.93 ± 1.13 s (mean ± S.D.) to rise from 10% to 90% of maximum, mimicking the encounter of a chemical in a fluid stream), followed by a fast decay phase (1.64 ± 1.04 s (mean ± S.D.) to fall from 90% to 10% of maximum, see example traces in **Figure 3C**, also see **Methods**).

**Figure 3.**
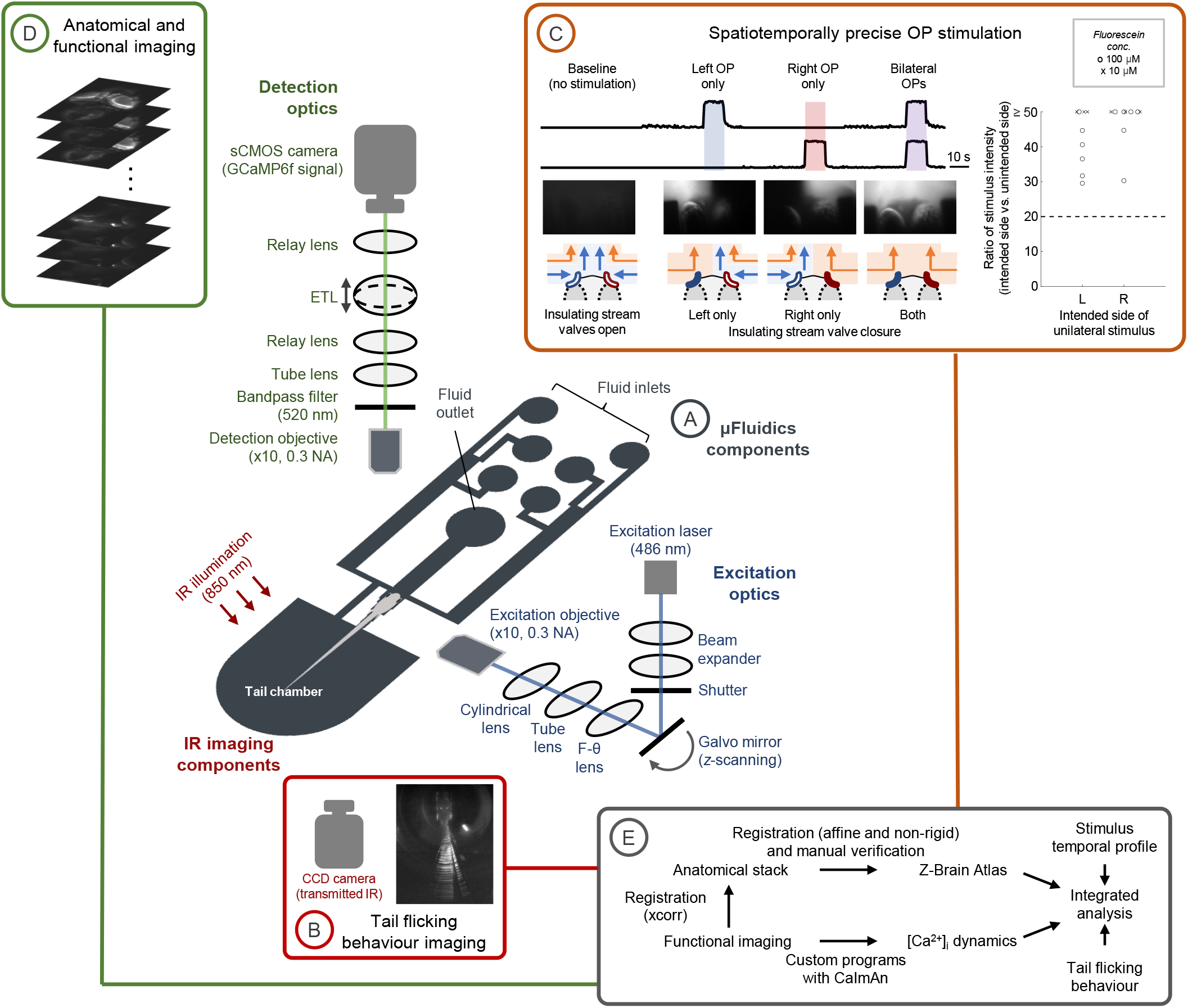
Integrated microfluidics-light sheet microscopy for larval zebrafish whole-brain imaging with spatiotemporally precise nasal stimulation. **(A)** In the system, a microfluidic module is integrated with a scanning light sheet microscope. The microfluidic module consists of a PDMS-based device with a tail chamber, a fish head and waist-trapping chamber, fluid inlets connected by tubings and pumps, gated by valves for fluid inflow control, and a tube-connected fluid outlet. Also see **Supplementary Figure 3A** for more detailed drawings and descriptions. **(B)** To monitor tail flicking behaviour, infrared (IR) illumination and imaging by a CCD camera permits simultaneous motor readout. **(C)** Spatiotemporally precise unilateral or bilateral nasal stimulation is achieved by shutting down the water stream(s) immediately in front of the OP(s), which otherwise insulate the cadaverine stream(s) (1 mM in water), on the left, right, or both sides (illustrated at the bottom row). 1 μM fluorescein in the cadaverine stream permits the visualization and monitoring of chemical delivery during functional imaging experiments (upper panels correspond to epifluorescence imaging for monitoring the fluid streams, with the readouts shown above). For clarity, the water streams further away to the chemical streams are not shown in this illustration. See **Supplementary Figure 3C** for more details. Right panel: Ratio of stimulus intensity (intended side vs. unintended side) in test trials (*n* = 9 trials from 3 larvae) using higher fluorescein concentrations (circles: 100 μM; crosses: 10 μM) in the cadaverine stream. All recorded ratios were > 20 (dashed line). **(D)** On the excitation path, a light sheet is formed by using a cylindrical lens and underfilling the back aperture of the excitation objective, with *z*-scanning achieved by a galvo mirror. On the detection arm of the optics, an electronically tunable lens (ETL) synchronized to the galvo mirror focuses the GCaMP6f signals from the different axial planes of the larval zebrafish brain onto the sCMOS camera. **(E)** The data analysis pipeline consists of custom-written programs for (i) regional identification in detailed anatomical stack by affine and non-rigid volumetric image registration to the Z-Brain Atlas as the common reference, (ii) calcium dynamics extraction with CaImAn, and regional identification by registration of functional imaging planes to the Z-Brain Atlas-registered anatomical stack, (iii) tail angle extraction, and (iv) integrated analysis with stimulus information (i.e., null, left OP (l-STIM), right OP (r-STIM), or bilateral OP (b-STIM) stimulation) and motor readout.

Using our integrated microfluidics-light sheet imaging system, we performed volumetric whole-brain calcium imaging in 5–6 d.p.f. larval zebrafish expressing the calcium reporter GCaMP6f in neurons^78^, spanning 29 planes with 7-μm intervals at 2 Hz (**Figure 3D**), while cadaverine (1 mM) was presented to unilateral or bilateral OP(s). A custom code-based workflow incorporating anatomical registration to the Z-Brain Atlas^79^, time-series image registration^80^, and calcium signal extraction^81^ was developed for integrated data analysis (**Figure 3E,** see **Methods**). Due to the orthogonal optical path configuration of light sheet imaging, a significant proportion of the forebrain regions between the eyes could not be imaged when a larval subject was placed at an upright orientation (**Supplementary Figures 3D and 3E**). Therefore, we performed imaging in two configurations that permitted the incoming excitation light sheet to illuminate the majority of the whole brain, including the forebrain regions (**Supplementary Figures 3F–I**): (1) tilting the larvae by up to 20° to the right (**Supplementary Figures 3F and 3G**), or (2) with surgical ablation of the right eye of the larvae 12–24 hours prior to imaging (**Supplementary Figures 3H and 3I**). Both approaches circumvented the occlusion of the forebrain from imaging (**Supplementary Figures 3G and 3I**). Trials of different stimulus conditions (null, left OP stimulation (l-STIM), right OP stimulation (r-STIM), or bilateral OP stimulation (b-STIM)) were presented in a randomized order.

We carried out experiments in 18 right eye-ablated and 6 tilted larvae with at least 3 occurrences of each stimulus condition. Among the right eye-ablated larvae, 50% (9 out of 18) exhibited tail flicking in the device, while none of the tilted larvae were behaviorally responsive. Comparing the tail flick events of the behaviorally responsive larvae across the stimulus conditions, we noted tendencies of higher tail flick frequency and angular velocity during bilateral cadaverine stimulation than control (i.e., trials with no stimulation) or unilateral stimulation (**Supplementary Figures 4A–E**). Compounded by the feasible number of repetitions, the stochastic nature of response to cadaverine, and possibly physical restraint and behavioral passivity of the larvae^82,83^, there were no statistically significant differences in these parameters across the conditions (**Supplementary Figures 4A–E**). Although the larvae were not as responsive as in the swimming arena, the system permitted us to study brainwide chemosensory representation, and identify the regional distribution of neurons that encode sensory stimulation-associated motor output.

### Brainwide neuronal activation patterns with olfactory placode stimulation and motor output

Along the early olfactory pathway of larval zebrafish, olfactory bulb (OB) neurons project to bilateral OBs and diverse forebrain regions^67,68^, but it is unclear whether the ipsilateral or the contralateral pathway is functionally dominant. Further down the pathways, it is also not known how unilateral or bilateral nasal inputs determine the propagation of neuronal activity and sensory representation. These uncertainties made it difficult to constrain models of whole-brain neural circuits mediating chemosensory processing and behaviors.

We first analyzed neuronal activity evoked by cadaverine under each stimulus condition. We identified sensory-encoding neurons, defined as those with the most significant mutual information (*I*_S_) between activity and cadaverine stimulus, with l-STIM, r-STIM or b-STIM (see **Methods**), from 9 stably imaged larvae (*n* = 6 behaviorally responsive, right eye-ablated and 3 tilted-imaged larvae with sufficiently stable recordings for high-quality across-trial registrations). We separated our analyses for animals exhibiting unbiased vs. biased neural responses to r-STIM (see **Methods** and **Supplementary Figures 4F–I** for classification and details). For unbiased animals (*n* = 4 larvae), *I*_S_ of individual neurons were directly further analyzed (**Figures 4A and 4B**). In a separate analysis, *I*_S_ of individual neurons from each animal were normalized to the mean *I*_S_ of its left, right or bilateral OBs during l-STIM, r-STIM or b-STIM, respectively (*I*_N_), to permit pooling and visualization across all larvae (*n* = 9) (**Figures 4C and 4D**).

**Figure 4.**
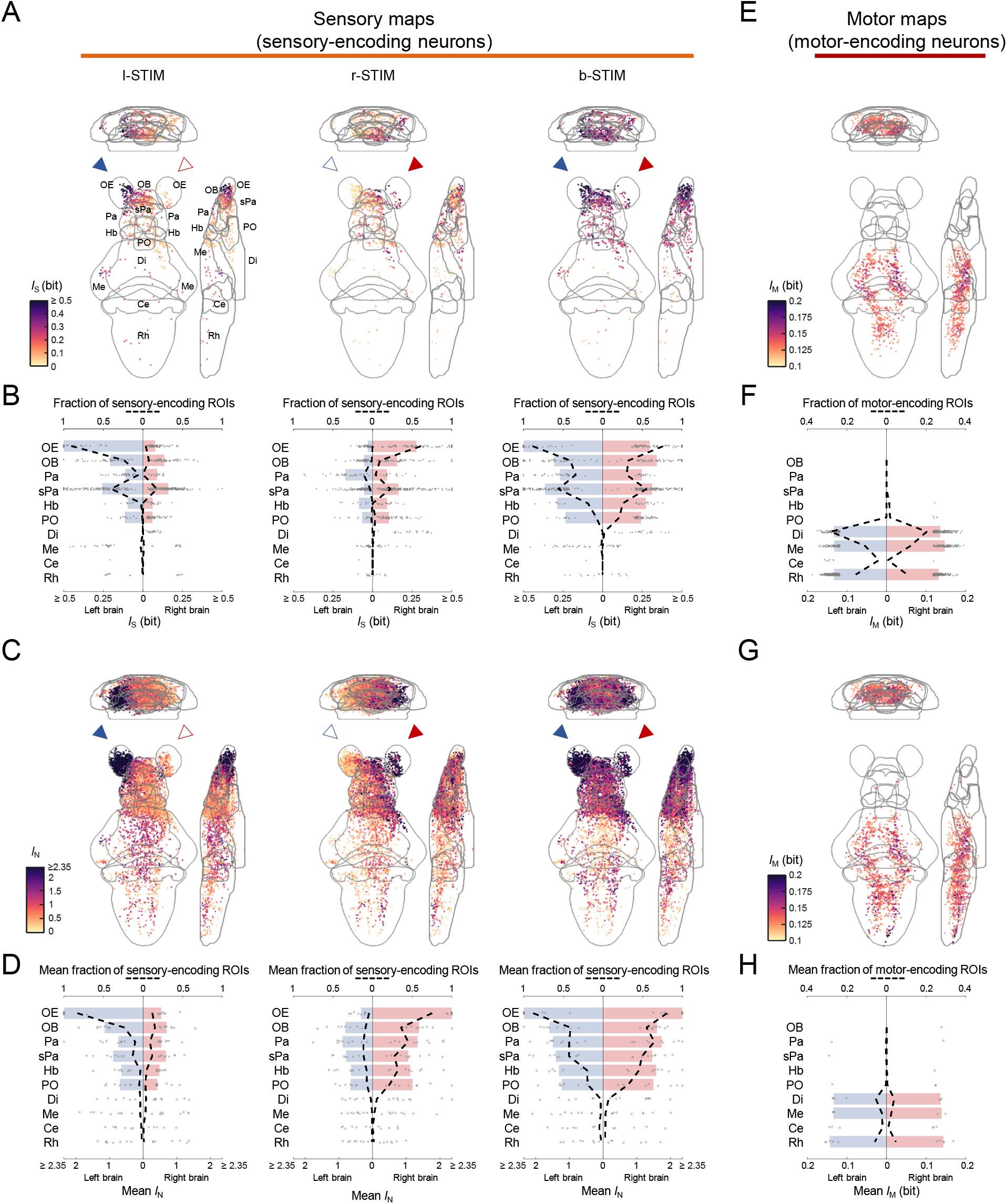
Brainwide sensory- and motor-encoding maps during spatiotemporally precise cadaverine stimulation. **(A)** Mean intensity projections (to coronal, transverse and sagittal planes) of the mutual information between the calcium signals of regions-of-interest (ROIs) and cadaverine stimulus profile of l-STIM (left panel), r-STIM (middle panel) or b-STIM (right panel) (*I*_S_ of sensory-encoding ROIs), from an example larval subject. Solid triangles mark the corresponding OP(s) stimulated. Abbreviations: OE, olfactory epithelium; OB, olfactory bulb; Pa, pallium; sPa, subpallium; Hb, habenula; PO, preoptic area; Di, diencephalon; Me, mesencephalon; Ce, cerebellum; Rh, rhombencephalon. **(B)** From the example fish in **(A)**, the distributions of *I*_S_ of sensory-encoding ROIs in the different brain regions with l-STIM (left panel), r-STIM (middle panel) or b-STIM (right panel). Regions with top six fractions of sensory-encoding ROIs with b-STIM are OE, OB, Pa, sPa, Hb and PO. Bars represent the medians of *I*_S_ of sensory-encoding ROIs in these regions. Dashed lines indicate fractions of sensory-encoding ROIs in all regions. **(C)** Mean intensity projections of the normalized mutual information between the calcium signals of ROIs and cadaverine stimulus profile of l-STIM (left panel), r-STIM (middle panel) or b-STIM (right panel) pooled across larvae (*n* = 9) (*I*_N_ of sensory-encoding ROIs). Solid triangles mark the corresponding OP(s) stimulated. **(D)** The distributions of mean *I*_N_ of sensory-encoding ROIs in the different brain regions during l-STIM (left panel), r-STIM (middle panel) or b-STIM (right panel) among the larvae (*n* = 9). Regions with top six mean fractions of sensory-encoding ROIs with b-STIM are OE, OB, Pa, sPa, Hb and PO. Bars represent the medians of the mean *I*_N_ of sensory-encoding ROIs in these regions. Dashed lines indicate mean fractions of sensory-encoding ROIs in the different regions among the larvae. **(E)** Mean intensity projections of mutual information between the calcium signals of ROIs and tail flick frequency (*I*_M_ of motor-encoding ROIs) from the same example larva in **(A)**. **(F)** The distributions of *I*_M_ of motor-encoding ROIs in the different brain regions, from the same example larva in **(A)**. Regions with top three fractions of motor-encoding ROIs are Di, Me and Rh. Bars represent the medians of *I*_M_ of motor-encoding ROIs in these regions. Dashed lines indicate fractions of motor-encoding ROIs in all regions. **(G)** Similar to **(E)** but pooled across larvae (*n* = 6). **(H)** Similar to **(F)** but pooled across larvae (*n* = 6), with bars representing the medians of mean *I*_M_ of motor-encoding ROIs, and dashed lines indicating mean fractions of motor-encoding ROIs, among the larvae.

We then examined whole-brain neuronal activity patterns (**Figures 4A–D, Supplementary Figures 4H and 4I**). Unilateral OP stimulation results in strong activation of olfactory sensory neurons (OSNs) in the ipsilateral OE. Along the olfactory pathway, we found robust bilateral neuronal activation as early as in the OBs. In the forebrain, neurons in the bilateral pallium (Pa, or dorsal telencephalon) and subpallium (sPa, or ventral telencephalon) both respond to unilateral OP stimulation. Although a known asymmetrical bulbo-habenular olfactory pathway projects selectively to the right habenula (Hb)^84^ which is preferentially activated by some odor cues^22,85^, in the majority of the larvae the left Hb neurons were similarly activated as the right counterpart by cadaverine. The preoptic area (PO), a hypothalamic region known to mediate nocifensive behaviors^63^, also generated bilateral neuronal activity during unilateral OP stimulation. There is progressively weaker sensory encoding along the rostral-caudal axis in these regions. Beyond the forebrain regions, there is only direct sensory encoding by small subsets of neurons in the diencephalon (Di), mesencephalon (Me), cerebellum (Ce) and rhombencephalon (Rh). Notably, during bilateral OP stimulation, symmetrical regional activation with an increase in the sensory information contents of individual neurons’ responses was observed. This points to a <u>binasal</u> input-dependent propagation of sensory-evoked neuronal activities.

From the behaviorally responsive larval zebrafish, we also calculated the mutual information between individual neuronal activity and tail flick movements recorded (*I*_M_) (see **Methods**). The motor-encoding neurons were mostly found in the caudal brain regions where fewer neurons are direct sensory-encoding (i.e., Di, Me and Rh), and only very few were found in the forebrain regions (**Figures 4E–H**). Collectively, we thus obtained the brainwide activation patterns underlying unilateral or bilateral nasal input processing, as well as the regional distributions of motor output-encoding units.

As control experiments, we performed imaging in 4 larval zebrafish in the upright configuration (with intact eyes). Among the optically accessible brain regions in these larvae, including partial forebrain regions and the caudal brain regions, we found similar relative levels of neuronal activation across brain regions in response to l-STIM, r-STIM or b-STIM (**Supplementary Figures 5A–D**). The regional trends of neuronal responses in 4 unilateral OP-ablated larvae with b-STIM were also similar to that evoked by l-STIM or r-STIM using the microfluidics system in larvae with intact OPs (**Supplementary Figures 5H and 5I**). From the larvae with intact OPs, we analysed neuronal activation in the trigeminal ganglia (gV) (**Supplementary Figures 5F and 5G**), as the terminal nerves also innervate the larval zebrafish OE^74,86^ and may participate in cadaverine chemosensation. However, sensory encoding by gV neurons were much weaker than those in the olfactory processing regions (**Supplementary Figure 5G**).

Since we observed that the nOP larvae evaded cadaverine similarly efficiently as the uOP larvae (**Figures 1D–H**), we performed imaging in 4 larval subjects with bilateral OPs ablated. In these larvae, unilateral or bilateral stimulation still elicited relatively strong neuronal responses in multiple forebrain areas (**Supplementary Figures 7A and 7B**). Given the geometric constraints of chemical stimulation in the microfluidic device, these were likely due to gustatory stimulation. This observation also suggested that binasal input integration properties, instead of the presence or absence of cadaverine-evoked activities, may underlie the differences in avoidance behavior in bOP vs. uOP, or nOP larvae.

### Binasal input integration in the early olfactory pathway and downstream brain regions for cadaverine sensing

We then investigated the rules of binasal sensory input integration for cadaverine sensing. Firstly, we characterized the propagation of information evoked from unilateral stimulation in terms of neuronal and regional selectivity to ipsilateral vs. contralateral sources. For the larvae with unbiased responses to unilateral stimulation (*n* = 4), we calculated the ipsilateral input selectivity of each neuron (defined as *I*_S_ipsi-STIM_/(*I*_S_ipsi-STIM_ + *I*_S_contra-STIM_), where *I*_S_ipsi-STIM_ and *I*_S_contra-STIM_ denote mutual information between calcium activity and ipsilateral or contralateral OP stimulus, respectively). In all regions except the OE, neurons fall within a wide distribution of input preference, ranging from being highly ipsilaterally selective to highly contralaterally selective (**Figures 5A and 5B**). In the OBs, where subsets of neurons respond to both ipsilateral and contralateral OP cadaverine stimulation, the overall higher ipsilateral selectivity inherited from the OE remains partially conserved (**Figures 5A–D**). The ipsilateral input bias becomes weaker in the Pa and the sPa, as well as in the other more caudal brain regions (**Figures 5A–C**). Therefore, while within each region a wide range of response selectivity is maintained, binasal input signals converge both early in the olfactory processing pathway and in the downstream regions. This indicates a brainwide, distributed mode of parallel input integration.

**Figure 5.**
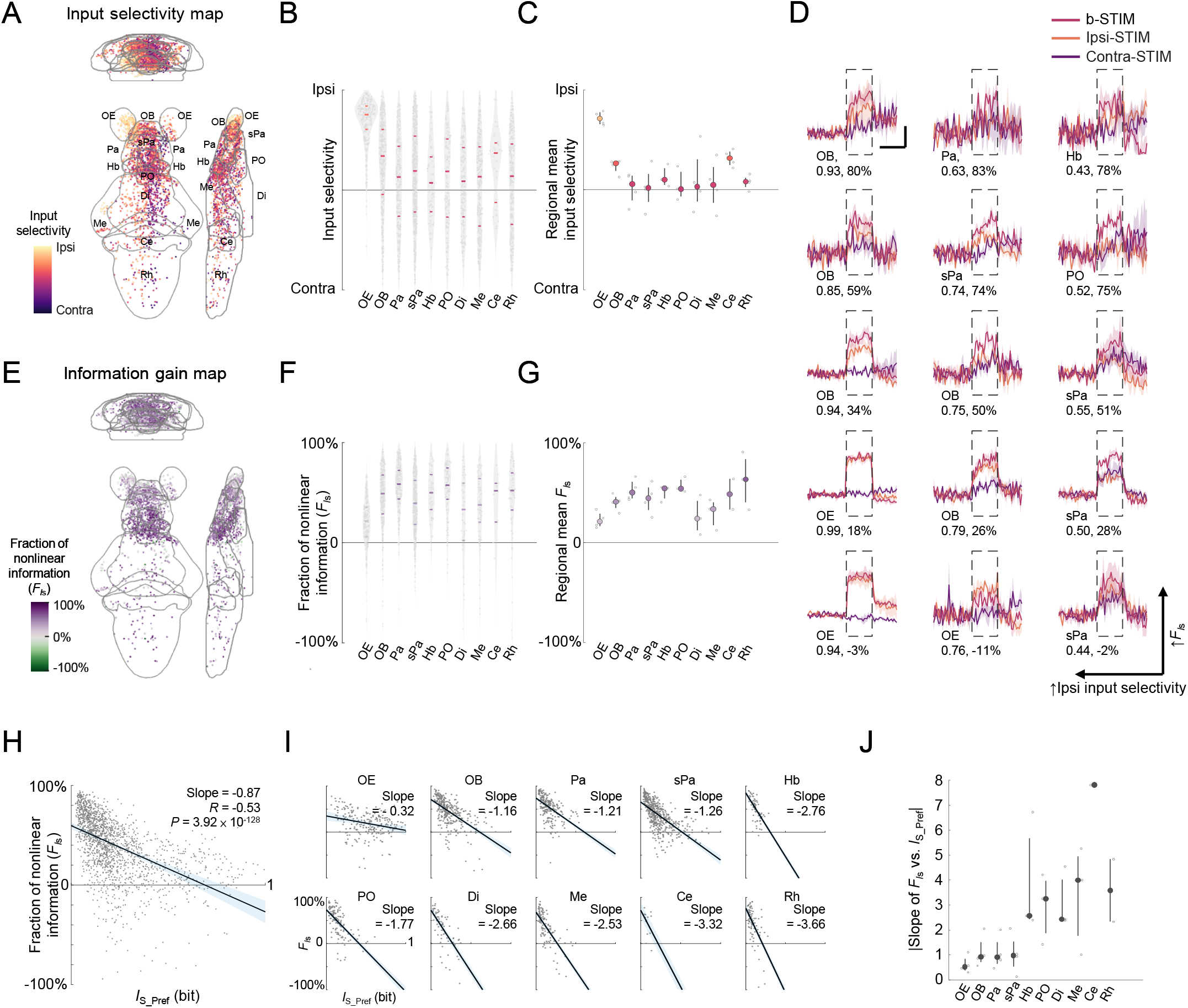
Bilateral nasal input integration properties underlying cadaverine sensing. **(A)** Mean intensity projection maps (to coronal, transverse and sagittal planes) of ipsilateral(Ipsi)-contralateral(Contra) input selectivity of sensory-encoding regions-of-interest (ROIs) (pooled across larvae, *n* = 4). Abbreviations of brain regions: same as in **Figure 4**. **(B)** Distributions of ipsilateral-contralateral input selectivity in the different brain regions. Horizontal lines: medians, 75 and 25 percentiles. **(C)** Regional means of ipsilateral-contralateral input selectivity, with each small dot representing the value from one larval zebrafish. Large dots, upper and lower limits of lines: medians, 75 and 25 percentiles, respectively. **(D)** Example trial-averaged responses to ipsilateral (ipsi-STIM, orange), contralateral (contra-STIM, violet) or bilateral (b-STIM, cherry) OP stimulation of individual ROIs from the designated brain regions with a range of ipsilateral-contralateral input selectivity (first number below each example, 1 represents total ipsilateral selectivity, while 0 represents total contralateral selectivity) and fraction of nonlinear information (i.e., fraction of information gained beyond linear summation, denoted by *F*_*I*s_) (second number below each example). Shadow shows SEM for each trace. Dashed rectangle indicates stimulus window. Scale bars: 10 seconds (horizontal) and 0.5 normalized *dF/F* (vertical). **(E)** Mean intensity projection maps of *F*_*I*s_ (pooled across larvae, *n* = 4). **(F)** Distributions of the *F*_*I*s_ of individual ROIs in the different brain regions. Horizontal lines: medians, 75 and 25 percentiles. **(G)** Regional means of *F*_*I*s_, with each small dot representing the value from one larval zebrafish. Large dots, upper and lower limits of lines: medians, 75 and 25 percentiles, respectively. **(H)***F*_*I*s_ vs. *I*_S_ during preferred unilateral OP stimulation by cadaverine (*I*_S_Pref_, defined as *max*(*I*_S_l-STIM_, *I*_S_r-STIM_), where *I*_S_l-STIM_ and *I*_S_r-STIM_ denote *I*_S_ with l-STIM and r-STIM, respectively) for ROIs pooled across regions and larvae. Black line and light blue shadow show line of best fit on linear regression with 95% confidence interval. Slope of the line of best fit, correlation coefficient (*R*) and *P*-values are shown. **(I)** Similar to **(H)** but with separate plot and linear regression performed for the ROIs of each brain region. Slopes of the lines of best fit are shown. **(J)** The absolute values of the slope of *F*_*I*s_ versus *I*_S_Pref_ for each brain region (i.e., |Slope of *F*_*I*s_ vs. *I*_S_Pref_|), with each small dot representing the value from one larval zebrafish. Large dots, upper and lower limits of lines: medians, 75 and 25 percentiles, respectively.

To examine the linearity of bilateral input integration, we next fitted generalized linear models (GLMs) to the responses of sensory-encoding neurons (**Supplementary Figure 6**) for all larvae (*n* = 9). We compared GLMs with kernels for l-STIM and r-STIM stimulus temporal profiles, and with or without an additional kernel accounting for left-right input interactions (i.e., b-STIM profile as an independent regressor) (**Supplementary Figures 6A and 6B**, also see **Methods**). We found that a GLM with the interaction kernel outperformed that without in terms of goodness-of-fit for the majority of neurons (**Supplementary Figure 6C**), indicating nonlinear interactions of the afferent input signals. Fitting GLMs with an additional kernel to account for the influence of activity history and calcium signal decay revealed similar results (**Supplementary Figures 6D–F**).

We hence further characterized the linearity of bilateral integration by determining the fraction of nonlinear information (i.e., fraction of information gained beyond linear summation, denoted by *F*_*I*s_) in larvae with unbiased responses to unilateral stimulation (*n* = 4). *F*_*I*s_ was defined as (*I*_S_b-STIM_ - *I*_S_u-STIM_sum_)/*I*_S_b-STIM_, where *I*_S_b-STIM_ denotes the mutual information of b-STIM-evoked responses and b-STIM stimulus profile, while *I*_S_u-STIM_sum_ denotes that of linearly summed unilaterally-evoked responses (see **Methods**). In principle, a neuron can have sublinear (*F*_*I*s_ < 0), near-linear (*F*_*I*s_ ≈ 0) or supralinear (*F*_*I*s_ > 0) gain of information with input summation. Interestingly, apart from observing a supralinear gain of information for the majority of neurons (**Figures 5D–G**), there is an increasing trend of *F*_*I*s_ (**Figures 5F and 5G**) as *I*_S_ values decrease across brain regions along the rostral-caudal axis (**Figures 4A–D**).

Additional analysis on right eye-ablated larvae with biased responses to r-STIM (*n* = 5) also revealed similar results on the trend of information gain across brain regions (**Supplementary Figures 7D–F**). Although only limited brain regions could be imaged in larvae (*n* = 4) in the upright configuration, we observed similar generally supralinear binasal input summation by neurons in the Di and Me (**Supplementary Figure 5E**). In contrast to the larval subjects with intact OPs, in the nOP larvae (*n* = 4) b-STIM did not increase the sensory information contents as supralinearly, with near-zero median *F_Is_* values in most of the forebrain regions (**Supplementary Figure 7C**). Nonlinear integration of bilateral stimuli therefore requires the integrity of the OPs, and cannot be accounted for by non-olfactory cadaverine chemosensation.

We speculated that the *F*_*I*s_ increase in the higher-order and caudal brain regions may be due to a larger relative information gain in neurons that weakly encode unilateral input, which could be more reliant on the convergence of the bilateral inputs. Consistent with the hypothesis, we found a strong inverse correlation between *F*_*I*s_ and the *I*_S_ with preferred OP stimulation (*I*_S_Pref_, defined as *max*(*I*_S_l-STIM_, *I*_S_r-STIM_), where *I*_S_l-STIM_ and *I*_S_r-STIM_ denote *I*_S_ of l-STIM and r-STIM conditions, respectively) (**Figure 5H**). Interestingly, in the higher and caudal brain regions, the relationship between *F*_*I*s_ and *I*_S_Pref_ becomes even stronger, whereby the magnitude of the negative slope of linear regression progressively increases (**Figures 5I and 5J**). This indicates that in these brain regions, given the same unit change in *I*_S_Pref_ there is a relatively larger change of *I*_S_b-STIM_ (**Figures 5I and 5J**). We noted the same phenomena in the larvae with biased responses to r-STIM (**Supplementary Figures 7G–I**). Synergistic input summation may therefore be especially important for sensory information representation by neurons that otherwise only weakly encode unilateral input in the downstream sensorimotor pathways. These include the caudal brain regions where the motor-encoding neurons are located (**Figures 4E–H**).

### Binasal input enhances sensorimotor unit responses in motor-encoding brain regions

Sensorimotor transformation requires sequential activation of neurons embedded in neural circuits specializing in roles from detection of sensory cues to driving motor outputs. The enhanced sensory representation by binasal inputs in brainwide regions should in principle be utilized by subsets of neurons mediating sensorimotor transformation and motor output. We thus examined how sensorimotor neurons may be modulated by unilateral vs. bilateral OP cadaverine stimulation in the subset of stably imaged larval subjects that were behaviorally responsive (*n* = 6). Sensorimotor neurons were defined as those that at least moderately encode both cadaverine stimulus information and motor output (see **Methods**). Rh had the largest number of sensorimotor neurons, followed by Di and then Me (**Figures 6A and 6B**). Most of the sensorimotor units exhibited supralinear sensory information gain with bilateral olfactory inputs, with *F*_*I*s_ higher than their sensory-encoding-only counterparts in the same regions (**Figure 6C**). This was also substantiated by examining their regionally averaged responses, as b-STIM resulted in larger stimulus-locked responses than unilateral stimulation (**Figures 6D and 6E**), which may result in stronger motor unit recruitment. Indeed, more motor-encoding neurons were identified when stimulus trials were included than only during baseline trials (**Figure 6A, *middle* and *right***). In the sensorimotor transformation process, the binasal input-enhanced sensory representation is therefore associated with a stronger drive to the downstream sensorimotor and motor-encoding neurons (**Figure 7**).

**Figure 6.**
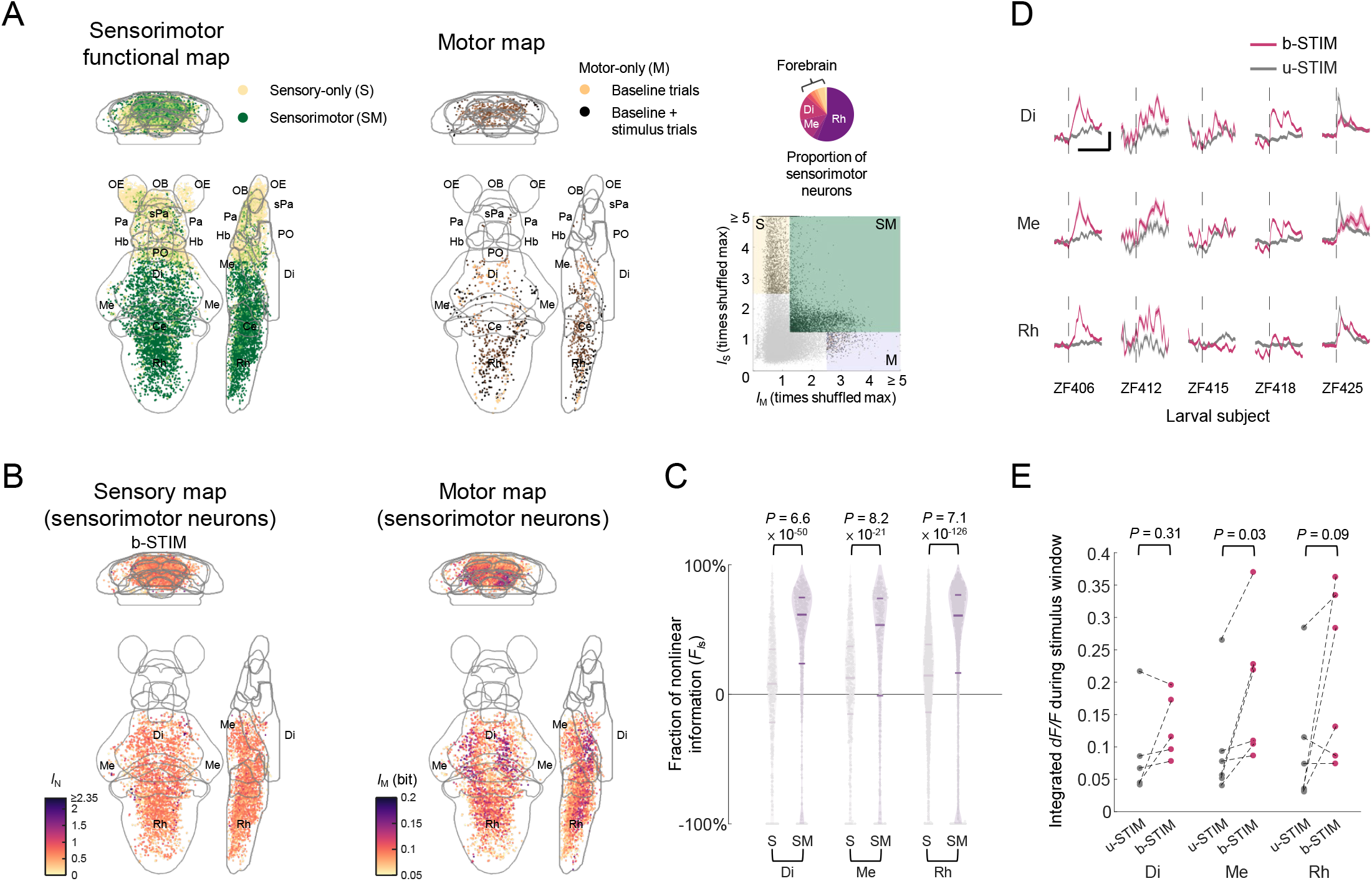
Binasal input-dependent activation of sensorimotor units in cadaverine sensing. **(A)** Left panel: Whole-brain functional maps (projections to coronal, transverse and sagittal planes) of sensory-only (S) and sensorimotor (SM) ROIs pooled across the behaviorally responsive and stably imaged larvae (*n* = 6). Middle panel: Whole-brain maps of baseline trial-motor-encoding (baseline trials) and non-baseline trial-motor-encoding (baseline + stimulus trials) ROIs. Right upper panel: Pie chart showing the distribution of sensorimotor ROIs in the different brain regions. Right lower panel: Scatter plot showing the distributions of the mutual information between the calcium signals of each ROI and cadaverine stimulus profile (maximum *I*_S_ of l-STIM, r-STIM and b-STIM conditions), and tail flick frequency (*I*_M_), both calculated as the number of times the maximum of the corresponding shuffled values (see **Methods**). Abbreviations of brain regions: same as in **Figure 4**. **(B)** Mean intensity projection maps of the mutual information between the calcium signals of sensorimotor ROIs and cadaverine stimulus profile of b-STIM (left panel, normalized mutual information *I*_N_) and tail flick frequency (right panel, mutual information *I*_M_), in Di, Me and Rh pooled across the same group of larvae. **(C)** Distributions of the fraction of nonlinear information (*F*_*I*s_) of sensory-only (S) and sensorimotor (SM) ROIs in Di, Me and Rh. Horizontal lines: medians, 75 and 25 percentiles. *P*-values: Wilcoxon rank-sum test. **(D)** Regionally averaged stimulus-locked sensorimotor neuronal responses by bilateral (b-STIM, cherry) and unilateral (u-STIM, grey) cadaverine stimulation from 5 larvae with sensorimotor ROIs identified in all three regions. Vertical dashed line indicates stimulus onset. Scale bars: 10 seconds (horizontal) and 0.5 normalized *dF/F* (vertical) from each larval subject. Shadows: SEM. **(E)** Integrated regionally averaged sensorimotor neuronal responses (i.e., summed *dF/F* over stimulus window) to b-STIM (cherry) and u-STIM (grey). Each pair of connected dots represents data from one larval zebrafish. *P*-values: Wilcoxon signed-rank test.

**Figure 7.**
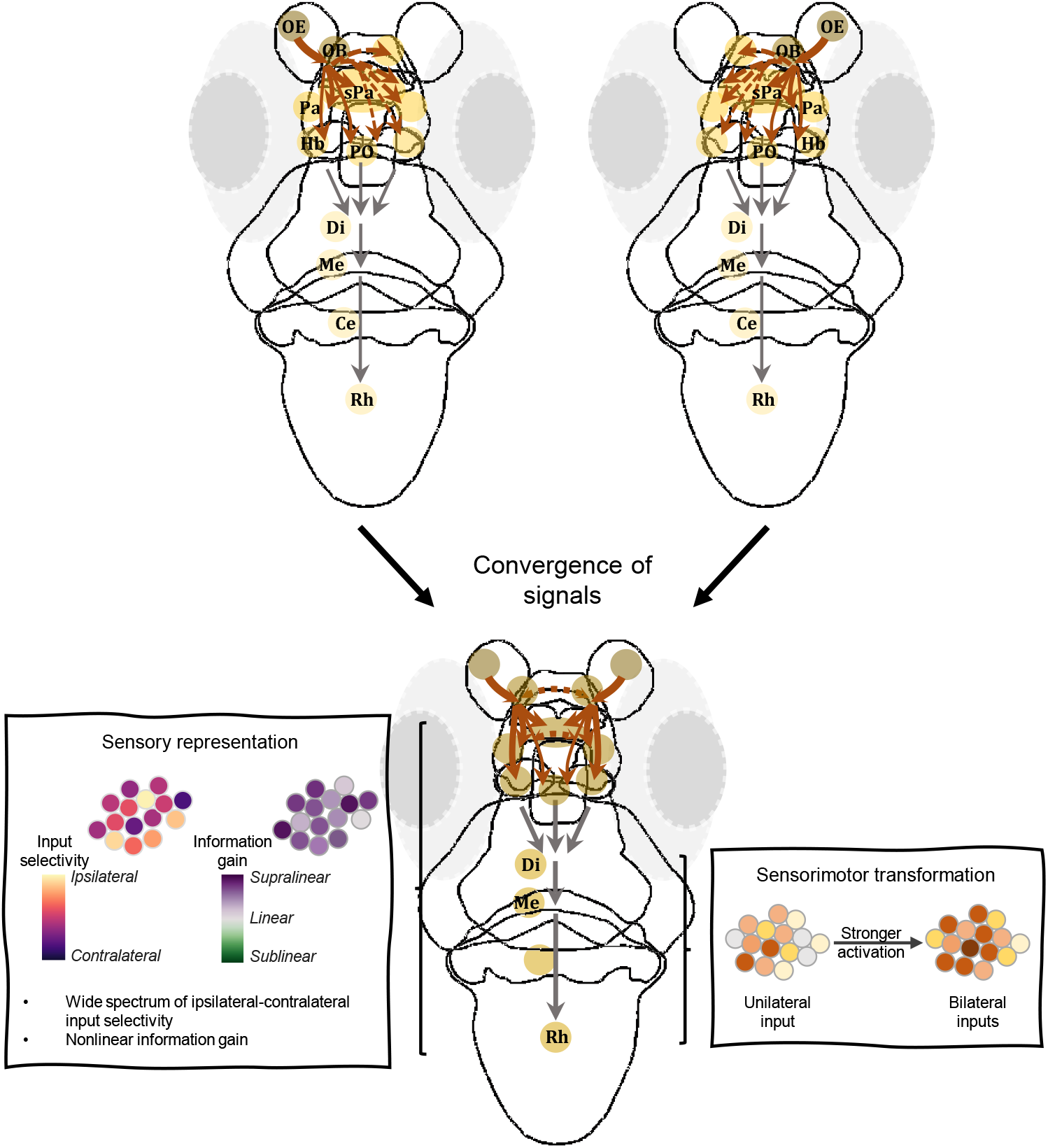
Illustration of brainwide convergence, integration and propagation of olfactory signals in cadaverine sensing. Upper panels: Each olfactory bulb receives inputs from the ipsilateral olfactory epithelium. Upon stimulation, the information is broadcasted to multiple forebrain and downstream brain regions, via ipsilateral and commissural projection pathways. With only unilateral input, the sensory information representation steeply becomes weaker along the rostral-caudal axis. Solid brown arrowed lines represent ipsilateral signal propagation, while dashed brown arrowed lines represent signal propagation to the contralateral side. Lighter colors of circles indicate regions with weaker sensory information representation in cadaverine sensing. Grey lines represent propagation of signals to downstream sensorimotor processing regions. Lower panel: Throughout the olfactory processing pathways, the diversity of stimulus selectivity is preserved. The convergence of signals from the bilateral inputs and their synergistic interactions results in a distributed sensory representation, whereby odor information is especially enhanced (relatively) in the higher-order and caudal brain regions. Meanwhile, this results in stronger activation of sensorimotor neurons. This could be the neural basis of the cadaverine-modulated turn behaviors that exhibit a dependence on binasal inputs.

## Discussion

Considerable progress has been made in circuit neuroscience in the past decade, in part owing to methodological advancements permitting large-scale cellular-resolution imaging of neuronal activities in diverse brain areas in various species, including zebrafish^60,75,76,87^, *Drosophila^88,89^*, and rodents^90–92^. Whole-brain imaging in larval zebrafish with several microscopic techniques, whereby the activities of as many as nearly 80% of all neurons could be imaged^60,75,76,87^, opened up the avenue for the study of brainwide circuit dynamics at cellular resolution in a vertebrate species^43–45^. It also permitted the assumption-free identification of brain regions mediating different behaviors. Taking advantage of the approach, in this work we developed microfluidics-based methodologies to empower the investigation of chemosensory-mediated behaviors and its neural basis in the larval zebrafish. These included a behavioral assay with a precisely defined and static chemical landscape, and an odorant delivery methodology with high spatiotemporal precisions for the interrogation of whole-brain neural representations. These tools were indispensable to reveal binasal input-dependent avoidance behavioral changes to cadaverine, as well as the underlying neural basis of parallel sensory input integration and subsequent sensorimotor transformation in the vertebrate model.

Cadaverine is an aquatic odor of ecological significance. In several species, diamines are detected by a family of trace amine-associated receptors (TAARs)^93^. The zebrafish genome has relatively many (109) TAAR genes^94^, compared with tetrapods. The adult zebrafish have been shown to express TAAR13c (a TAAR) in a subset of ciliated OSNs that mediate their cadaverine avoidance behavior^95,96^. Intriguingly, in a previous study, puff-application of cadaverine did not elicit immediate escape response in zebrafish larvae^74^, in apparent contradiction to the danger-signifying and toxic nature of the chemical upon prolonged exposure. By allowing larvae to navigate in a nature-mimicking arena with precisely defined and continuous chemical-carrying laminar flows, our assay captured the behavioral responses upon naturalistic encounters of cadaverine. Repeated sampling of a large number of cadaverine zone entry events and escape journeys enabled us to reveal adaptive behavioral changes that occur in a statistical manner.

Although still in the early developmental stage, the larval zebrafish has a rich behavioral repertoire with complex dynamical properties. Its spontaneous locomotor activity is characterized by discrete swim bouts with correlated turn-direction bias in successive bouts^97,98^. The adaptive nature of locomotion is exemplified by at least thirteen swimming patterns used in different combinations depending on the behavioral contexts^97^. For instance, light modulates locomotion, and zebrafish larvae predominantly swim in slow bouts in darkness^97^. In our experiments, to avoid the potential confounding effects of light or other visual stimuli, and to isolate the effects of odor stimuli, we carried out the avoidance behavioral assays in the absence of visible light. Our data supports that under such context, the larval zebrafish employs a combined strategy of klinokinesis and angular orthokinesis, without directional bias, to efficiently evade cadaverine. It appears to be a sensible strategy, given the small spatial separation of the OPs at this stage likely renders chemical gradient determination by inter-OP comparison sensitive to random fluctuations or noises in sampling. It is yet to be addressed whether with further development, zebrafish may learn to compare odor arrival times or intensity differences across OPs for more efficient odorant localization like larger mammals^1,14–16^.

Interestingly, we found that larvae with unilateral or bilateral OPs ablated could still avoid cadaverine, albeit less efficiently than their normal counterparts. Larval zebrafish cadaverine avoidance therefore encompasses a nasal input-independent component, and its optimization is binasal input-dependent. Consistently, while an increasing number of intact nasal inputs increases angular orthokinesis in a graded manner, unilateral nasal input is insufficient to enhance klinokinesis beyond that driven by OP-independent cadaverine sensing. These suggest that angular orthokinesis is principally nasal input-driven, whereas klinokinesis consists of both strictly binasal input-dependent and independent components. As the terminal nerves of the trigeminal system were likely ablated together with the OP(s) in our experiments, the nasal input-independent component of klinokinesis may be mediated by gustatory cadaverine detection. It remains unclear what may account for the bilateral input-dependent adaptation of turn duration (and thereby turn magnitude) with continuous cadaverine exposure. Adaptation of sensorimotor transformation, glial and neuromodulatory system-mediated behavioral futility with successive unsuccessful attempts^82,83^, and downstream motor output adaptations (e.g., at neuromuscular or muscular levels) could in theory all contribute to it. Nonetheless, its presence suggests that once a larval subject fails to evade in the initial swim bouts, subsequent attempts may become less efficient. Therefore, it is behaviorally even more pertinent to perform the first few turns with sufficiently large angle and displacement upon entering a noxious or toxic chemical-carrying stream.

To study the neural basis of binasal input integration that underlies the adaptive behavioral changes to cadaverine sensing, the prerequisite is a method for spatiotemporally precise delivery of odor stimuli to unilateral or bilateral OPs. This requires partitioning of the chemical delivery fluid streams at ten-micrometer precision. Careful direction of fluid flows around a larva subject is therefore needed, which is achievable by microfluidics techniques with suitable fluid channel geometry design and fluidic pressure control. In our setup, we harnessed the laminar property of fluid flow at low Reynolds number, such that distinct pairs of fluid streams are separately directed to the OPs from the sides and converge at the midline with minimal mixing. With a spatial separation of ~100 μm between the two OPs, the maximal tolerable deviation of midline without contacting the opposite OPs is approximately ± 50 μm, well above the typical length scale of microfluidic flow (~5 μm). In order to minimize the delay of fluid changes which is intrinsically variable (subject to momentary local fluidic resistance), we aimed to switch fluids as close to the larva as possible. We therefore used insulating water streams between the OPs and the cadaverine streams, which can be rapidly switched off for repeated trials of temporally reproducible odor delivery.

With obvious ecological significance, the olfactory system and its associated pathways develop early in the zebrafish. Within 7 d.p.f., the OB projection pathways that target diverse forebrain regions are already formed, which include extensive inter-hemispheric projections^67,68^. Although we observed a small overall bias of neuronal responses to ipsilateral OP cadaverine stimulation, many neurons in each of the brain regions are equally driven by the left and the right stimuli. This suggests that while the ipsilateral projection pathways may be slightly stronger, neither the ipsilateral nor the contralateral pathways functionally dominate. The functional roles of commissural pathways likely vary depending on the specific computational goals to achieve^29,30,42^. In the adult zebrafish, the interbulbar pathways improve the detection of a reproductive pheromone especially when there are co-existing confounding odor cues^30^. In cadaverine sensing, we found that binasal input convergence confers a generally supralinear sensory information gain in the OBs and downstream regions. The interbulbar and the other commissural olfactory pathways are therefore already functional at the larval stage, and mediate overall excitatory or facilitatory interactions, as we observed predominantly positive response modulation by the odor stimuli. Our results also suggest that they likely serve to improve the signal-to-noise ratio of odor detection, and enhance subsequent sensorimotor transformation to mediate cadaverine avoidance. In theory, other advantages offered by dual sensory channels may also include (i) expanded dynamic range of signal detection as each sensory organ is physically limited by certain maximum neuronal packing capacity, (ii) preserved possibility of stereo-comparison as the spatial separation between the pair of organs increases with growth, and (iii) as a fail-safe design when one input channel is damaged.

On the other hand, despite the early signal convergence in the olfactory bulbs, a wide range of response selectivity to each of the nasal inputs is maintained in the higher-order olfactory processing centers and the caudal brain regions. This distributed representation preserves information on sensory cue laterality. If conserved with further development, the design may provide the neural substrate for juvenile or adult zebrafish to potentially utilize the information for guiding directed turns, thereby further optimizing avoidance behavior. Although in trials with only unilateral OP stimulation intended, we could not ensure that there was absolutely no cadaverine spillover to the contralateral OP, the presence of unintended contralateral OP stimulation may lead to an underestimation of both the unilateral input selectivity and supralinearity of input summation (by exaggerating unilateral nasal input-evoked activities).

In our experiments, it was not possible to completely dissociate behavioural and neuronal activity modulations due to concurrent gustatory stimulation, or fully rule out any involvement of the trigeminal system. Yet, the supralinearity of input summation vanished without intact nasal inputs, which correlated with weakened klinokinesis and angular orthokinesis, highlighting the unique importance of binasal cadaverine detection and the olfactory system in optimizing avoidance behavior. Although calcium signals recorded with GCaMP6 imaging could be nonlinear with respect to the instantaneous firing rate, over a reasonable firing rate range of typical sensory-evoked activities, action potential number-calcium relationship tends to be relatively linear^78,99,100^. We therefore propose that the nonlinearity of calcium signals with respect to firing rate would not fully account for the supralinear summation we observed, and we hypothesize that the underlying mechanism depends on cellular and circuit-based input summation nonlinearity. Testing this hypothesis requires electrophysiological recordings of both suprathreshold and subthreshold activities from individual neurons, while chemical stimulation is presented to unilateral or bilateral OPs.

Originally intended to improve the optical accessibility of forebrain regions, we unexpectedly observed biased responses to the ipsilateral nasal input in the majority of larvae imaged after right eye ablation. The absence of right eye stimulation appeared to potentiate the responses to ipsilateral olfactory stimulus and decrease that to the contralateral side. There are several possible explanations for the crossmodal response modulation. A potential mechanism is lateralized sensory compensation^101^, whereby removal of the right eye may have led to a heightened odor sensitivity of the ipsilateral OE, while a yet-to-be characterized inhibitory pathway may suppress that of the contralateral counterpart. To avoid the phenomenon and eliminate the need to image in a tilted configuration, future works will require improved microfluidic device design that can simultaneously incorporate more fluid inlets and accommodate an additional narrow light sheet from the front to illuminate the forebrain, while the light sheet entering from the side can then conveniently avoid the eyes^102^.

Finally, the microfluidics-based behavioral, chemical delivery and imaging platforms we developed can be readily adapted for studying chemosensory behaviors in other organisms of similar or smaller length scales, such as bacteria, *C. elegans* (see related works ^103,104^), *Drosophila* larvae and adults. In fact, other fluid properties (e.g., temperature, viscosity) can also be precisely controlled. Convenient incorporation of crossmodal stimuli allows the study of their combined effects on the behaviors and neural responses of animal subjects. The high-precision delivery system is also extensible to accommodate an elaborate array of chemicals varying in species, concentrations and valence, to interrogate the corresponding neural representations in the larval zebrafish. In this work, by delivering congruent odors to the OPs, we showed how bilateral input integration may permit odor perceptual enhancement. On the other hand, by delivering incongruent odors with opposite valence to the OP(s), one may investigate how competing circuits implement the prioritization of more ecologically important sensory cues. These will gain us deeper insights into the architecture of chemosensory information processing in vertebrates.

## Acknowledgements

We thank Tom Mrsic-Flogel and Florian Engert for helpful comments on the manuscript; Jan Schnupp and Vincent Cheung for insightful discussions; Florian Engert for provision of the Tg(elavl3:GCaMP6f) zebrafish. We thank Becky Yung, Florence Yau, Anki Miu and Rebecca Chau for administrative support to the project. This work was in part funded by a Croucher Innovation Award from the Croucher Foundation (CIA20CU01) and a Faculty Innovation Award (FIA2017/B/01) from the Faculty of Medicine, the Chinese University of Hong Kong (CUHK) (H.K.); the Gerald Choa Neuroscience Centre, the Margaret K. L. Cheung Research Centre for Parkinsonism Management, Faculty of Medicine, CUHK (V.C.T.M. and H.K.); the Collaborative Research Fund (C6027-19GF) and the Area of Excellence Scheme (AoE/M-604/16) of the University Grants Committee of Hong Kong (H.K.).

## Author Contributions

S.K.H.S. constructed experimental setups, carried out experiments and analyzed data. D.C.W.C. contributed to image processing. R.C.H.C. and J.L. assisted imaging experiments. Z.L. assisted two-photon ablation experiments. H.M.L. contributed to experimental protocol testing. K.K.Y.W., C.H.J.C. and V.C.T.M. contributed to data interpretation. Y.H. advised data analysis and interpretation. S.K.H.S., D.C.W.C. and H.K. wrote the manuscript with input from all authors.

## Declaration of Interests

The authors declare no competing interests.

## STAR ★ METHODS

### KEY RESOURCES TABLE

| REAGENT or RESOURCE | SOURCE | IDENTIFIER |
| --- | --- | --- |
| Chemicals |  |  |
| Cadaverine | Sigma | D22606-5G |
| Experimental Models: Organisms/Strains |  |  |
| Tg(elavl3:GCaMP6f) zebrafish larvae | Engert lab (Harvard) | ZFIN_ZDB-ALT-150916-3 |
| Software and Algorithms |  |  |
| Custom codes written in MATLAB R2018b | This paper | https://www.mathworks.com/products/matlab |
| AutoCAD 2018 | Autodesk | https://www.autodesk.com/products/autocad/overview?support=ADVANCED |
| StreamPix | Norpix | https://www.norpix.com/products/streampix/streampix.php |
| CFDTool | Precise Simulation | https://github.com/precise-simulation/cfdtool/raw/master/CFDTool.mlappinstall |
| Variational Stationary Noise Remover | Ref <sup>105</sup> | https://github.com/sattarab/image-quality-tools/tree/master/metrix_mux/metrix/vsnr |
| NoRMCorrE | Ref <sup>80</sup> | https://github.com/flatironinstitute/NoRMCorre |
| CaImAn | Ref <sup>81</sup> | https://github.com/flatironinstitute/CaImAn-MATLAB |
| Fiji - ImageJ | Ref <sup>106</sup> | http://fiji.sc |

### RESOURCE AVAILABILITY

#### Lead Contact

Further information and requests for resources and reagents should be directed to the lead contact, Ho Ko.

#### Materials Availability

The AutoCAD files of the microfluidic devices developed in this study are available upon reasonable request to the lead contact.

#### Data and Code Availability

Preprocessed data necessary to replicate these results will be deposited to G-Node, and the MATLAB code for data analysis will be made available on GitHub upon the acceptance of the paper.

### EXPERIMENTAL MODEL AND SUBJECT DETAILS

All experimental procedures were approved in advance by the Animal Research Ethical Committee of the Chinese University of Hong Kong (CUHK) and were carried out in accordance with the Guide for the Care and Use of Laboratory Animals. For all experiments reported in this study, we used 5–7 days post fertilization (d.p.f.) larvae carrying the *nacre* mutation. Larvae were raised and maintained under 14-h light:10-h dark cycles at 28°C. The larvae used for the bilateral OP-intact control and cadaverine groups in the navigation behavioral assay were bred from a pair of heterozygous Tg(elavl3:GCaMP6f) adult zebrafish without screening for GCaMP6f expression. For the unilateral OP-ablated groups, the larvae used in the assays had confirmed GCaMP6f expression (by examination under an epifluorescence microscope) to allow structural visualization and two-photon OP ablation.

### METHOD DETAILS

#### Microfluidic arena for chemosensory behavioral assays

The microfluidic device for the avoidance navigation assay was designed in AutoCAD 2018 (Autodesk, USA) and consists of three 1.5 mm-thick laser-engraved poly(methyl methacrylate) (PMMA) layers. The layers were vertically aligned and fused by a chloroform solution applied to the edges of the sheets. This created a closed rectangular arena (60 mm (L) × 30 mm (W) × 1.5 mm (H)) connected to three fluid inlets and two fluid outlets during assays (**Figure 1A**). An additional inlet was used for loading larvae and sealed prior to assays. The inlets and outlets are 200 μm wide, prohibiting larvae from exiting the arena through these channels. Syringe needles (Terumo, USA) were sealed at the top layers with epoxy adhesive paste (Devcon, USA) to connect the inlet and outlet channels to fluid-filled syringes and waste collection bottle, respectively, via poly(tetrafluoroethylene) (PTFE) tubes (Cole Parmer, USA). The microfluidic device was then air-dried for one day to allow evaporation of the residual chloroform and epoxy adhesive paste. The device was rinsed twice with water before use for behavioral assays.

To create a static chemical zone, three fluid streams (two water streams, and one chemical stream with 1 mM cadaverine dissolved in water (pH = 9.0) were formed in the arena by propelling the respective fluids through the inlet channels at 220 μL/min using syringe pumps (LSP02-2A, Longerpump, Beijing) (**Figure 1A**). The outlets were connected to open waste collection bottles. The Reynolds number (*Re*) of fluid flow in the arena is given by *Re* = *ρνD*_H_/*μ*, where *ρ* is the fluid density, *ν* is the fluid flow velocity, *D*_H_ is the hydraulic diameter of the device, and *μ* is the fluid dynamic viscosity. Given the rectangular cross-sectional dimensions of the arena (i.e., 1.5 mm × 60 mm), the average volumetric flow rate was 220 μL min^-1^ × 3 inlets = 660 μL min^-1^, which equals 11 mm^3^ s^-1^. Hence the average flow velocity = 11 mm^3^ s^-1^ / (1.5 mm × 60 mm) = 1.22 × 10^-4^ m s^-1^. *D*_H_ for the rectangular channel is given by 2*ab*/(*a + b*) = 9 × 10^-5^ m, where *a* and *b* are the dimensions of the rectangular cross section (i.e., 1.5 mm and 60 mm). Substituting the *D*_H_ found and the density (996 kg m^-3^) and dynamic viscosity (8.32 × 10^-4^ m^2^ s^-1^) of water at 28°C, the Reynolds number was found to be *Re* = 0.013. As *Re* << 2000, the streams were laminar with negligible mixing^107^ and static fluid zone borders were formed (**Figure 1A, Supplementary Figure 1B**). The static-border property of the chemical zone during the entire 2-hour assay period was validated by infusing an IR dye (IR 806, Sigma, USA) into the rightmost stream of the device in the presence of larvae (**Supplementary Figure 1B**). Sharpness of the border was quantified by the relative IR dye intensity (normalized to mean 0 outside, and 1 within the stimulus zone) of 15 line profiles perpendicular to the border evenly distributed from the top to the bottom end (**Supplementary Figure 1C**).

During experiments, the device was levelled and the outflow rates of the two outlets were identical. The two-dimensional (2D) flow velocity profile (**Supplementary Figure 1A**) was obtained by computational fluid dynamics simulation using the CFDTool toolbox (Precision Simulation) in MATLAB (R2018b, MathWorks, USA), assuming incompressible Newtonian fluid flow and with a grid size of 1.4 × 10^-4^ m. The geometry and boundary conditions were set according to the aforementioned device parameters.

#### Chemosensory avoidance navigation and survival assays

During a chemosensory avoidance navigation assay, three to six larvae aged 5–7 days d.p.f. were placed in a swimming arena and allowed to acclimate for 15 minutes. Swimming and navigational behaviors were then recorded by a high-speed camera at 240 fps (Mako G-030B PoE, Allied Vision, Germany) under 850-nm IR illumination (ANGX-1000-CH1-24V, TMS, Malaysia) for 2 hours in the absence of visible light. Fluid temperature was maintained at ~28°C throughout the assay.

For the control group, all three fluid streams consisted of water. For the cadaverine zone avoidance assay groups, the rightmost stream contained 1 mM cadaverine (Sigma, USA) dissolved in water (**Figure 1A**), which was freshly prepared prior to each experiment due to the evaporative nature of cadaverine. For the unilateral or bilateral OP-ablated groups, larvae aged 4–5 d.p.f. were first individually immobilized in 1.5% low-melting agarose (Sigma, USA), then underwent two-photon ablation of the OP(s) which was performed under a customized two-photon microscope (Scientifica, UK) with a Ti:sapphire femtosecond laser (Spectra-Physics, USA) tuned to 830 nm. The unilateral or bilateral OP-ablated larvae were released from agarose immediately after ablation and allowed to recover in a water tank for at least 24 hours prior to experiments. Before the behavioral assays, the OPs were embedded in agarose again and imaged under the two-photon microscope to ensure successful ablation (see the left panels of **Figures 1C and 1D**). The numbers of assays, larvae and border-crossing event sample sizes for the four experimental groups are as follows: bilateral OP-intact larvae (control group without cadaverine zone, i.e., all water zones): 6, 24, 211; bilateral OP-intact larvae (assay group with cadaverine zone): 8, 32, 251; left OP-intact larvae (assay group with cadaverine zone): 4, 15, 71; right OP-intact larvae (assay group with cadaverine zone): 4, 15, 155; bilateral OP-ablated (i.e., null OP-intact) larvae: 7, 23, 96.

For the survival assays, larvae were individually put in a device filled with 1 mM cadaverine dissolved in water without flow. Videos of the larvae were recorded at 10 fps, and stimulated with gentle taps to the device every 15 minutes. The assay was terminated when the larva became unresponsive and death was confirmed by the absence of heartbeats and circulation under microscopic examination. No larvae survived continuous cadaverine exposure more than 165 minutes. A control group of larvae underwent the same assay with devices filled with water for 180 minutes.

#### Microfluidic device for precise olfactory placode stimulation, behavioral and brainwide imaging

Studying bilateral nasal input integration in larval zebrafish requires fine spatiotemporal control of chemical-carrying fluid flow around the larval subject. On the other hand, stable cellular-resolution neuronal imaging of partially restrained larval zebrafish wholly immersed in continuous fluid medium during OP-specific stimulus presentation must be attained. The PDMS-based microfluidic devices developed to achieve these goals (**Figure 3A and Supplementary Figure 3A**) were designed in AutoCAD 2018 (Autodesk, USA). Slight customizations were made to the larva head and waist-trapping chambers to allow the fitting of either an upright-oriented, right-tilted or right eye-ablated larva. The microfluidic device was fabricated by customized photolithography techniques to incorporate a flat and smooth sidewall for excitation light sheet entry, and a thin glass ceiling for emitted fluorescence detection path (**Supplementary Figure 3A**). Briefly, a 2D design was printed on a soda lime mask (Supermask, Shenzhen). Negative photoresist SU-8 2150 (Microchem, USA) was spin-coated on a 4-inch silicon wafer (Semiconductor Wafer, Taiwan) to 500 μm-thick. The device pattern was then transferred to the photoresist with UV exposure (OAI, USA), followed by post-exposure bake and etching to produce a master mold. Mixed PDMS (Dow Corning, USA) and curing agent (at 10:1 w/w) was poured into the salinized mold to a thickness of 5 mm, with a perpendicular glass slide held upright on the mold to produce a smooth vertical surface upon curing (**Supplementary Figure 3A**). After solidification at 100°C for 60 minutes, it was detached from the mold, bonded and sealed onto a microscopic cover slide (as thin glass ceiling) by plasma treatment. The devices were rinsed with water twice before use in experiments.

We derived a fluidic stream control solution to achieve the necessary spatial (**Supplementary Figure 3B**) and temporal precisions of olfactory placode stimulation (**Supplementary Figure 3C**). On each side anterior to the larva, there were sandwiched laminar fluid streams (water-cadaverine-water) which converged at the midline. The cadaverine stream consisted of 1 mM cadaverine dissolved in water, with 1 μM sodium fluorescein for the visualization of chemical delivery in stimulus-response mapping experiments (whereas additional characterization of devices with a larva in situ were carried out with 10 and 100 μM sodium fluorescein). Each fluid stream was carried by PTFE tubes, controlled by an solenoid valve (LHDA 0533115H, Lee Company, USA) and driven by a syringe pump. The posterior water streams on each side insulates the OPs from contacting the corresponding cadaverine streams (**Figure 3C, Supplementary Figure 3C, *left***). Closure of the valve of the insulating water stream on one side removes the water barrier to the chemical and enables precise chemical stimulation of the ipsilateral OP (**Figure 3C, Supplementary Figure 3C, *middle***). Unintended stimulation of the contralateral OP is prevented by the presence of opposing flow on the contralateral side (**Figure 3C, Supplementary Figure 3C, *middle***). Stimulation is turned off by closure of the chemical stream valves (**Supplementary Figure 3C, *right***). Bilateral stimulation is achieved by closure of the insulating water stream valves on both sides (**Figure 3C**) and terminated by closure of the chemical stream valves. A side channel connecting the chemical delivery microchannels and the tail chamber was incorporated to buffer pressure changes in the fluidic environment, and minimize mechanical disturbances to the larval subject. Flow rate of each stream were maintained at 88 μL/min with slight adjustments for each larva to compensate for the differences in position in the device, ensuring precise convergence of the opposing fluid streams at the midpoint between the OPs.

#### Light sheet microscope for whole-brain calcium imaging

We used a custom-built light sheet microscope for cellular-resolution whole-brain imaging (**Figure 3**). A 486 nm-centered blue gaussian laser (DL-488-050-0, Crystalaser, USA) was used as the excitation light source. The laser beam was resized to 0.6 mm in diameter (1/*e*^2^) by a pair of telescopic lenses (LB1757-A, LB1596-A, Thorlabs, USA), which then passed through a scanning system, followed by a cylindrical lens (LJ1695RM-A, Thorlabs, USA) that focused the horizontal dimension of the parallel beam onto the back focal plane of an air excitation objective (Nikon Plan Fluorite, ×10, N.A. 0.3, 16 mm WD). This expanded the laser horizontally to form a light sheet. The scanning system consisting of a galvanometric mirror (GVS211/M, Thorlabs, USA), a F-theta lens (S4LFT0061/065, Sill Optics, Germany) and a tube lens (TTL200-B, Thorlabs, USA) was used to scan the light sheet vertically and linearly over a range of 210 μm. The F-theta and tube lenses also expanded the beam 3.31 times, resulting in a beam diameter of 2 mm (1/9^th^ of the objective back aperture) and effective excitation N.A. of 0.0332. Thus, a thin excitation sheet (thickness: 7.33 μm (1/*e*^2^), Rayleigh length: ~300 μm) was generated.

Along the detection path, a bandpass green filter (525 ± 25 nm) was placed after an air detection objective (Nikon Plan Fluorite, ×10, N.A. 0.3, 16 mm WD) to block the blue excitation light. An electrically tunable lens (EL-10-30-Ci-Vis-LD, Optotune, Switzerland) in between a pair of relay lenses (LA1509-A, Thorlabs, USA) was linearly driven by a lens controller (TR-CL-180, Gardasoft, UK) and synchronized with the light sheet scanner to achieve rapid focusing of different image planes onto the sensor of a sCMOS camera (Zyla 5.5, Andor, UK). During an experiment, to correct for axial drift, a reference plane was calibrated with respect to the initial measurement at the beginning of each trial. All control units were synchronized using a multifunctional I/O device (PCIe-6323, National Instruments, USA).

#### Simultaneous whole-brain calcium imaging and tail flicking behavior recording

Eighteen 5–6 d.p.f. right eye-ablated and four right-tilted zebrafish larvae underwent simultaneous behavioral and whole-brain calcium imaging experiments. For the right eye-ablated group, 4.5–5 d.p.f. larvae were lightly anesthetized in 0.016% MS-222 and the right eyes were gently removed using surgical forceps under a dissection microscope. Larvae were returned to warmed Ringer’s solution and allowed to recover for 12–24 hours before imaging experiments. Additional control experiments were performed in four bilateral OP-intact (upright configuration), four bilateral OP-ablated, two left OP-ablated and two right OP-ablated larvae (right-tilted configuration) (see **Supplementary Table 1** for larvae used for each plots in the figures). For all experiments, larval subjects were only picked if they exhibited normal spontaneous swimming in the tank. Both right before and after imaging experiments, the larvae’s heartbeat and response to tactile stimuli were examined. All larvae included in the datasets exhibited normal heart rate and responded to tactile stimuli.

In an imaging experiment, a single larva was loaded and fitted into the trapping chamber of the PDMS-based microfluidic device using a syringe pump and monitored under a surgical microscope. The larvae inlet was then sealed and the larva was allowed to acclimate for 15 minutes. The axial focus of the light sheet was adjusted to be centered at the brain, correcting for small variations in side wall width and larva position inside the device. Prior to functional imaging, a detailed anatomical stack of the larval zebrafish brain spanning 138 imaging planes at 2-μm intervals was taken.

Whole-brain calcium imaging spanning 29 planes at 7-μm intervals was performed at 2 Hz volumetric rate, with each trial lasting for 50 or 70 seconds, and ~90-second inter-trial intervals. Trials consisted of 30-second baseline, 10- or 25-second stimulus presentation, and 10- or 15-second post-stimulus epochs. In between the trials, a resting interval was followed by axial drift correction. Each larva underwent 3–4 trials of null stimulus condition recordings and 3–6 trials of each cadaverine stimulus condition (i.e., left, right or bilateral OP stimulation). During each trial, after a 30-second baseline recording period, the first (water) and the second (cadaverine) stream in front of the OP(s) were switched off sequentially with 10- or 25-second gap to allow the second (cadaverine) followed by the third stream (water) to contact the OP(s), resulting in the initiation and termination of nasal stimulation, respectively (**Supplementary Figure 3C**). Recording continued for 10 or 15 seconds after stimulus cessation. Simultaneous tail flicking behavior was imaged at 200 fps under IR illumination by a CCD camera (F032B, Pike, Germany).

### QUANTIFICATION AND STATISTICAL ANALYSIS

#### Navigational behavior analysis

Individual frames of the navigation behavioral tracking videos were first registered for translation, background-subtracted, and contrast-adjusted. The heading orientation (10° resolution / accuracy) and *xy*-coordinates (0.1 mm resolution / accuracy) of every larva in each frame were extracted using a semi-automated template matching-based tracking program custom-written in MATLAB. All tracking results were manually verified on a frame-by-frame basis and corrected when necessary. When larvae came in close proximity (≤ 4 mm) to either the arena boundary or with each other, the trajectories were excluded from analysis of turn behaviors to rule out potential social or mechanical cue-associated movements. A cadaverine-water border was defined with the aid of an IR dye (IR 806, Sigma, USA) in a separate assay (**Supplementary Figure 1B**). A 1-mm margin along the border was incorporated in analysis, to account for the minimal diffusion and mixing. For each assay, the footprints of larvae in the arena (**Figures 1C and 1D**) were quantified by summing the head center coordinate occupancy of each pixel by any larva over time and normalized with respect to number of larvae and assay time. The proportions of time spent in the stimulus zone vs. mirror water zone were then quantified by summing values within the respective zones and divided by that over the whole arena. The associated standard errors of mean across assays were also calculated. Larval zebrafish head orientation changes with angular velocity ≥ 1.2 °ms^-1^ or linear displacements with velocity ≥ 12 μm ms^-1^ were registered as swim bouts. The swim bout event detection results were manually verified and corrected when necessary (by S.K.H.S.). Cadaverine zone border-crossing events were identified when swim bout trajectories intersected the cadaverine-water border from the water zone. Subsequent escape trajectories were isolated and analysed to compare swim bout kinematic parameters and escape efficiency for the different experimental groups.

#### Analysis of tail flicking behavior

Each frame of tail flicking behavioral imaging was background-subtracted and the tail tip-waist angle was extracted by a custom MATLAB program. The tail tip-waist angle was defined as the angle between the long axis of the larval subject body and a straight line joining the tail tip and the waist. The extracted angles were manually verified on a frame-by-frame basis and used for the calculations of tail flick event frequency and angular velocity.

#### Image processing and calcium signal extraction

Detailed larval zebrafish brain anatomical stacks were registered to the Z-Brain Atlas^79^ using affine transformation followed by non-rigid image registration using custom scripts written in MATLAB. Functional imaging planes were matched to the corresponding anatomical stack by maximization of pixel intensity cross-correlation and manually verified. Stripe artefacts in the anatomical stacks and functional imaging frames were removed using the Variational Stationary Noise Remover (VSNR) algorithm^105^. Functional imaging frames were motion-corrected using the NoRMCorrE algorithm^80^. Trials with blurred frames due to in-frame drifts were discarded. Regions of interest (ROIs) corresponding to individual neurons were then extracted using the CaImAn package^81^. Anatomical landmarks and the regional identity of each ROI were verified by manual inspection (by S.K.H.S.). The stimulus temporal profiles of l-STIM or r-STIM were obtained as the average pixel intensity of a manually defined ROI located immediately anterior to the olfactory epithelium on each side and within the nasal cavities. In trials included for analysis with only unilateral OP stimulation intended, absence of or minimal cadaverine spillover to the contralateral side was confirmed for each experimental trial by intended vs. unintended stimulation side peak fluorescence intensity changes from background (*dF/F*) near 20:1 or above. Trials with seizure-like brain activity and/or unsatisfactory image registration were also excluded from analysis. For each larva, 2 – 4 repetitions or trials per stimulus condition were included for final analysis (except one larva from the group with unbiased response which had only one l-STIM trial).

#### Identification of sensory-encoding, motor-encoding, and sensorimotor neurons

The first 10.5 second (21 imaging frames) of the imaging data of each trial were excluded from analysis to avoid the inclusion of transient activity evoked by the onset of light sheet illumination. The minimum fluorescence intensity in the subsequent 4.5 seconds (9 imaging frames) of each ROI was used as the baseline fluorescence (*F*). The *dF/F* signals of remaining imaging frames were calculated as the calcium responses of each ROI. The left (*C*_L,*t*_) and right (*C*_R,*t*_) cadaverine stimulus profiles were extracted using the abovementioned ROIs directly in front of the respective OPs and normalized to the [0, 1] range. The cadaverine stimulus profiles used in bilateral OP-stimulated trials (*C*_L+R,*t*_) were obtained by averaging the left and right stimulus profiles and normalized to the [0, 1] range. We calculated the mutual information (*I*_S_) between the calcium responses of each ROI and cadaverine stimulus profiles under each stimulus condition by a method based on kernel density estimation of the probability density functions of variables^108^. Each ROI thus had three *I*_S_ values, namely *I*_S_l-STIM_, *I*_S_r-STIM_, and *I*_S_b-STIM_, corresponding to *I*_S_ values between evoked responses and *C*_L,*t*_, C_R,*t*_ and *C*_L+R,*t*_, respectively. To estimate the expected distributions of *I*_S_l-STIM_, *I*_S_r-STIM_, and *I*_S_b-STIM_ due to randomness (shuffled *I*_S_), we randomly shuffled the *dF/F* signals of each ROI (i.e., disrupting their temporal structures) and calculated another set of shuffled values of *I*_S_l-STIM_, *I*_S_r-STIM_, and *I*_S_b-STIM_ (with each ROI contributing one value to each of the distributions). We defined sensory-encoding neurons to be those that most significantly encode cadaverine stimulus, with at least one *I*_S_ value > 2.5 times the maximum value of the pooled shuffled *I*_S_l-STIM_, *I*_S_r-STIM_, and *I*_S_b-STIM_ distributions. To calculate normalized *I*_S_ (i.e., *I*_N_), the *I*_S_ of individual ROIs of a given larva were normalized to the mean *I*_S_ of its left, right or bilateral OBs, during l-STIM, r-STIM or b-STIM, respectively.

For each sensory-encoding ROI, we calculated its right input selectivity, defined as *I*_S_r-STIM_/(*I*_S_l-STIM_ + *I*_S_r-STIM_), where *I*_S_l-STIM_ and *I*_S_r-STIM_ denote mutual information between calcium responses and left or right OP stimulus profiles, respectively. We classified the larval subjects to have biased responses if the mean right input selectivity of all sensory-encoding ROIs (except the OE) deviated from a balanced-response value (i.e., 0.5) by more than 0.15 (i.e., > 0.65 or < 0.35). Those with mean right input selectivity ≥ 0.35 and ≤ 0.65 were considered generally unbiased. In 5 out of 6 stably imaged right eye-ablated and behaviorally responsive larvae, we obtained r-STIM-biased responses in the majority of brain regions (except left olfactory epithelium (OE), see **Supplementary Figures 4F and 4G**). This phenomenon was not observed for the remaining right eye-ablated larva and 3 tilted-imaged larvae (**Figures 4A and 4B, Supplementary Figures 4H and 4I**). See **Supplementary Table 1** for details of larval subjects / datasets in each of the plots with pooled data across multiple larvae.

We defined motor output to be the proportion of frames (200 fps) with active tail flicking in each 0.5-s time bins. The calcium response-motor output mutual information (*I*_M_) was calculated with the same kernel density estimator-based method using data from all stimulus conditions. Shuffled *I*_M_ values were calculated using temporally shuffled motor output traces (with each ROI contributing one value to the distribution). Motor-encoding neurons (**Figures 4 and 6**) were defined as those that most significantly encode motor output, with *I*_M_ > 2.5 times the maximum values of the shuffled distributions. Motor-encoding neurons were further classified into baseline trial motor-encoding or non-baseline trial motor-encoding units (i.e., during null stimulus trials), defined with an *I*_M_ during baseline trials > 2.5 times the maximum values of the corresponding shuffled distributions, and the complementary set of remaining motor-encoding neurons, respectively.

Sensorimotor neurons (**Figure 6**) were defined as those with at least moderate encoding of both sensory (at least one *I*_S_ > 1.25 times maximum of the shuffled *I*_S_ values) and motor output (*I*_M_ > 1.25 times maximum of the shuffled *I*_M_ values) information. Sensory-only neurons (**Figure 6**) were classified based on strong sensory encoding (at least one *I*_S_ > 2.5 times maximum of the shuffled *I*_S_ values) and lack of significant motor encoding (*I*_M_ ≤ 1.25 times maximum the shuffled *I*_M_ values) (i.e., equivalent to sensory-encoding neurons defined as above with an additional constraint on the lack of motor encoding). Motor-only neurons (**Figure 6**) were classified based on strong motor encoding (*I*_M_ > 2.5 times maximum of the shuffled *I*_M_ values) and lack of significant sensory encoding (all *I*_S_ ≤ 1.25 times maximum of the shuffled *I*_S_ values) (i.e., equivalent to motor-encoding neurons defined as above but with an additional constraint on the lack of sensory encoding).

For the response traces shown (**Figures 5D, Supplementary Figures 6B and 6E**), normalized *dF/F* traces were calculated by dividing the *dF/F* values by the maximum of the trial-averaged traces of all stimulus conditions, to facilitate visualization and comparison of each ROI’s or region’s averaged responses across stimulus conditions.

#### Analysis of input selectivity and fraction of nonlinear information with bilateral input integration

Ipsilateral input selectivity for sensory-encoding neurons was defined as *I*_S_ipsi-STIM_/(*I*_S_ipsi-STIM_ + *I*_S_contra-STIM_), where *I*_S_ipsi-STIM_ and *I*_S_contra-STIM_ denote mutual information between calcium responses and ipsilateral or contralateral OP stimulus profiles, respectively. For a ROI in the left brain, *I*_S_ipsi-STIM_ and *I*_S_contra-STIM_ were equivalent to *I*_S_l-STIM_ and *I*_S_r-STIM_, respectively. Likewise, for a ROI in the right brain, *I*_S_ipsi-STIM_ and *I*_S_contra-STIM_ were equivalent to *I*_S_r-STIM_ and *I*_S_l-STIM_, respectively.

The fraction of nonlinear information (*F*_*I*s_) of each ROI was defined as (*I*_S_b-STIM_ - *I*_S_u-STIM_sum_)/*I*_S_b-STIM_, where *I*_S_u-STIM_sum_ denotes the mutual information between linearly summed unilateral OP stimulation-evoked responses and *C*_L+R,*t*_ (see above). The linearly summed unilateral OP stimulation-evoked responses were computed by adding the trial-averaged non-preferred OP (i.e., the OP with lower *I*_S_ upon stimulation) stimulation-evoked response trace to each preferred OP stimulation-evoked response trace. For this analysis, either ROIs with *I*_S_b-STIM_ > 2.5 times the maximum value of shuffled *I*_S_b-STIM_ (**Figure 5, Supplementary Figures 5 and 7**) or sensorimotor neurons (**Figure 6**) were included.

#### Generalized linear model (GLM) fitting

For the calcium responses of sensory-encoding neurons, we fitted Lasso regularised and cross-validated generalized linear models (GLMs) with custom scripts written in MATLAB. The regressors in the GLMs included the left (*C*_L,*t*_) and right (*C*_R,*t*_) cadaverine stimulus profiles, with or without (i) an activity history-dependence regressor, and (ii) a bilateral input interaction regressor (*C*_L∩R,*t*_, defined as the averaged left and right stimulus profile with values outside the time windows of bilateral stimulation set to zero, see **Supplementary Figures 6A and 6D**). In the simplest GLM (Model I), the calcium response at time *t* of a given ROI *R_t_* = *R*_0_ + *C*_L,*t*_ * *f*_L_ + *C*_R,*t*_ * *f*_R_, where * denotes convolution, *R*_0_ is a DC term, and *f*_L_ and *f*_R_ are the kernels fitted for the left and right stimulus regressors, respectively. To add a bilateral input interaction regressor (Model II), the model became *R_t_* = *R*_0_ + *C*_L,*t*_ * *f*_L_ + *C*_R,*t*_ * *f*_R_ + *C*_L∩R,*t*_ * *f*_L∩R_, where *f*_L∩R_ is the kernel fitted for the bilateral input interaction regressor. With the addition of an activity history-dependence term, the fitted GLMs became (Model III) *R_t_* = *R*_0_ + *C*_L,*t*_ * *f*_L_ + *C*_R,*t*_ * *f*_R_ + *R’_t_* * *f*_h_, and (Model IV) *R_t_* = *R*_0_ + *C*_L,*t*_ * *f*_L_ + *C*_R,*t*_ * *f*_R_ + *C*_L∩R,*t*_ * *f*_L∩R_ + *R’_t_* * *f*_h_, where *R’_t_* denotes activity history in the preceding 4 frames (i.e., [*R*_*t*-4_, *R*_*t*-3_ *R*_*t*-2_, *R*_*t*-1_]), and *f*_h_ the fitted kernel for the activity history (i.e., modeling up to 2-s preceding activity history-dependence). The kernel sizes for the l-STIM, r-STIM or b-STIM regressors was 10 (i.e., convolving with stimulus signals in a preceding 5-s window). Coefficients of best fit were determined as those that minimized cross-validation errors. For response prediction using the fitted GLMs, we used an identity link function. Root-mean-square error (RMSE) of each model for each ROI were calculated and compared accordingly.

**Supplementary Table 1.**
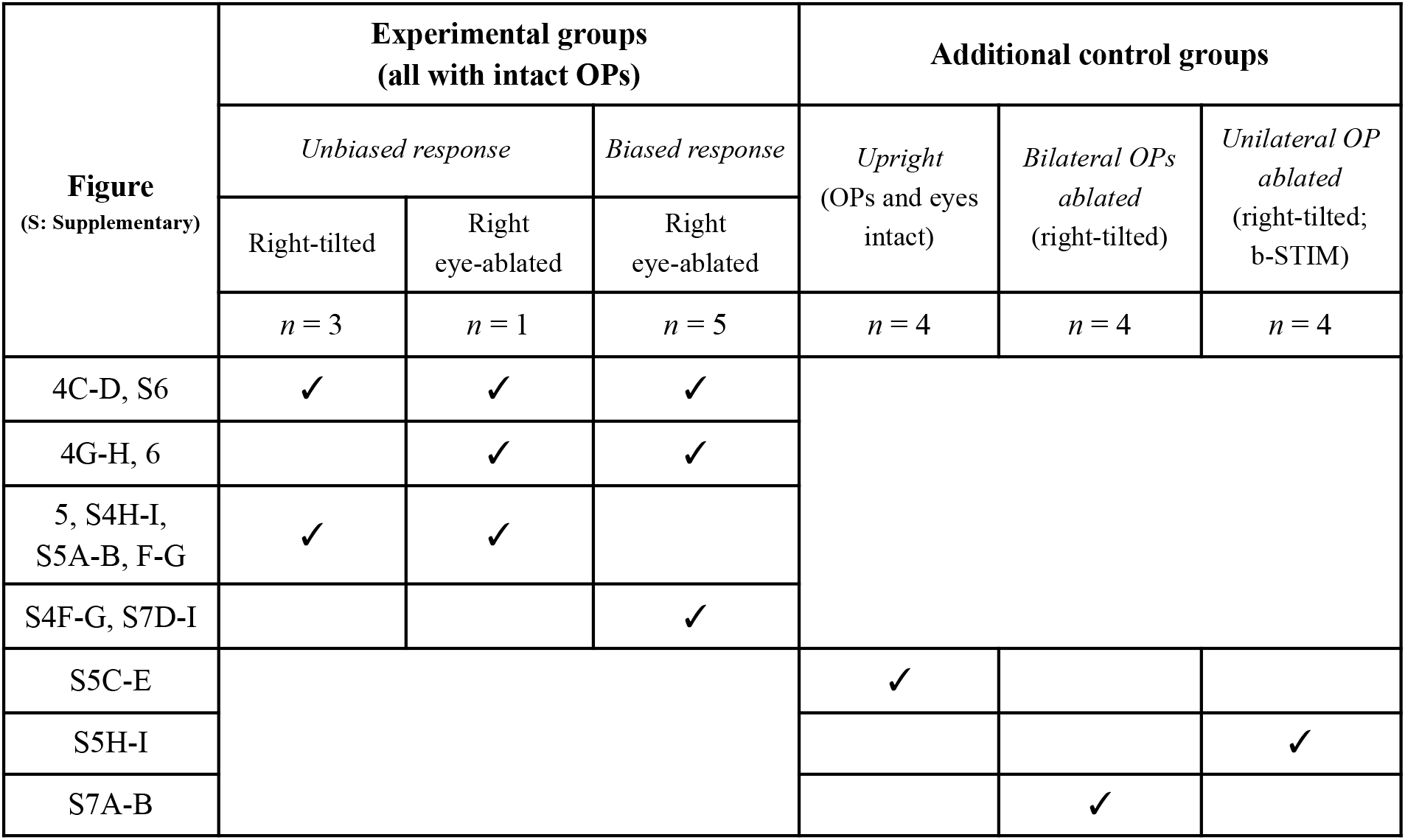

---

**Supplementary Figure 1.**
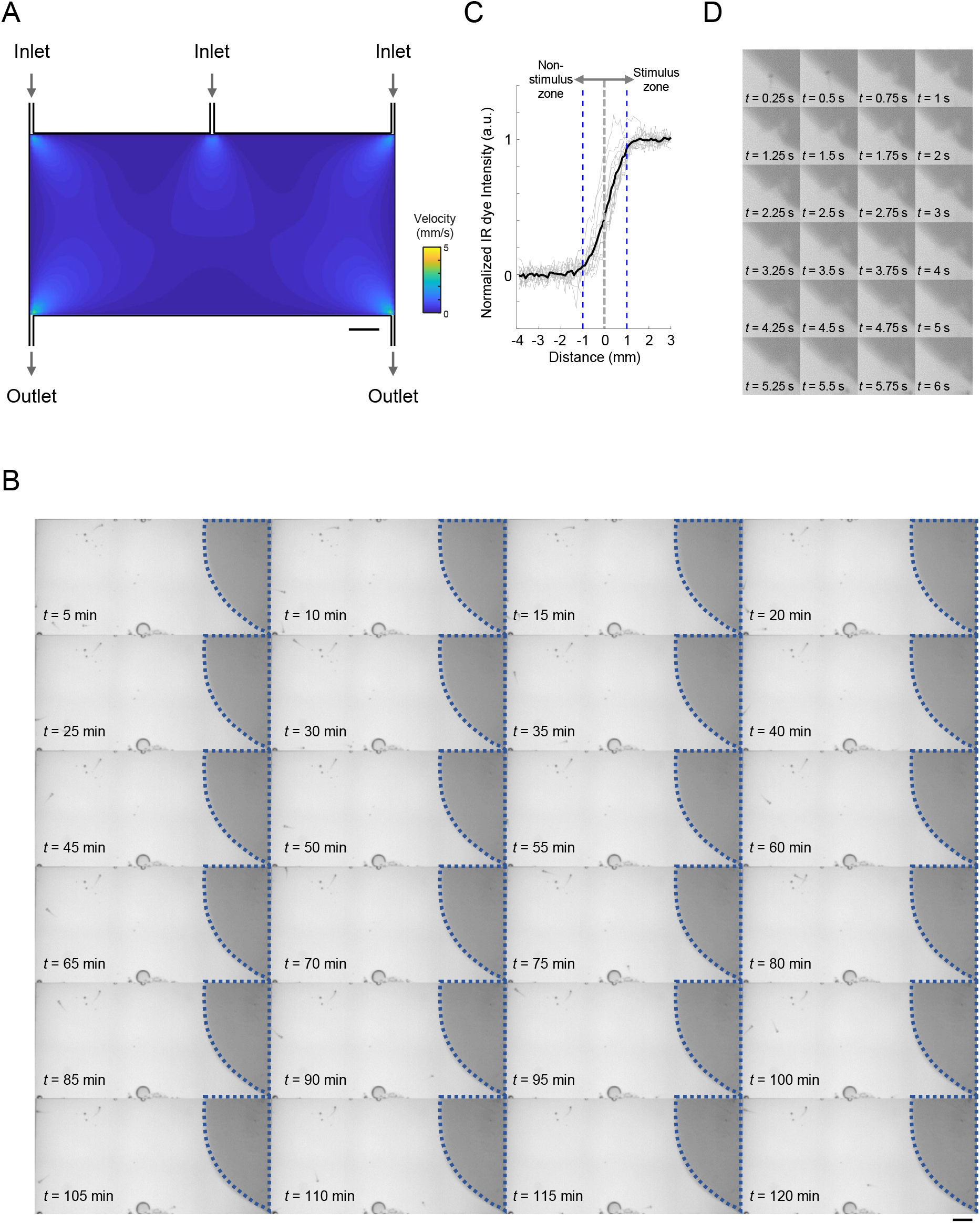
Maintenance of a static chemical zone in the fluidics-based avoidance behavioral assay. **(A)** Simulated fluid velocity profile in the swimming arena for avoidance behavioral assay. **(B)** Time-lapsed images showing the chemical zone border in the presence of larval zebrafish with an infrared (IR) dye flowing in via the rightmost fluid inlet in a swimming arena identical to that used for avoidance behavioral assay. Dotted lines in each image outline the same border. Scale bars in **(A)** and **(B)**: 0.5 cm. **(C)** Individual line profiles (grey) of IR dye intensity along normal vector at different spatial locations along the chemical border (see **Methods**) and their mean (black). Positive and negative values indicate distances from the border further into and away from the stimulus side, respectively. Blue dotted lines mark ± 1 mm from the border. **(D)** Time-lapsed images showing transient border disturbance and restoration in a larval zebrafish border-crossing event.

**Supplementary Figure 2.**
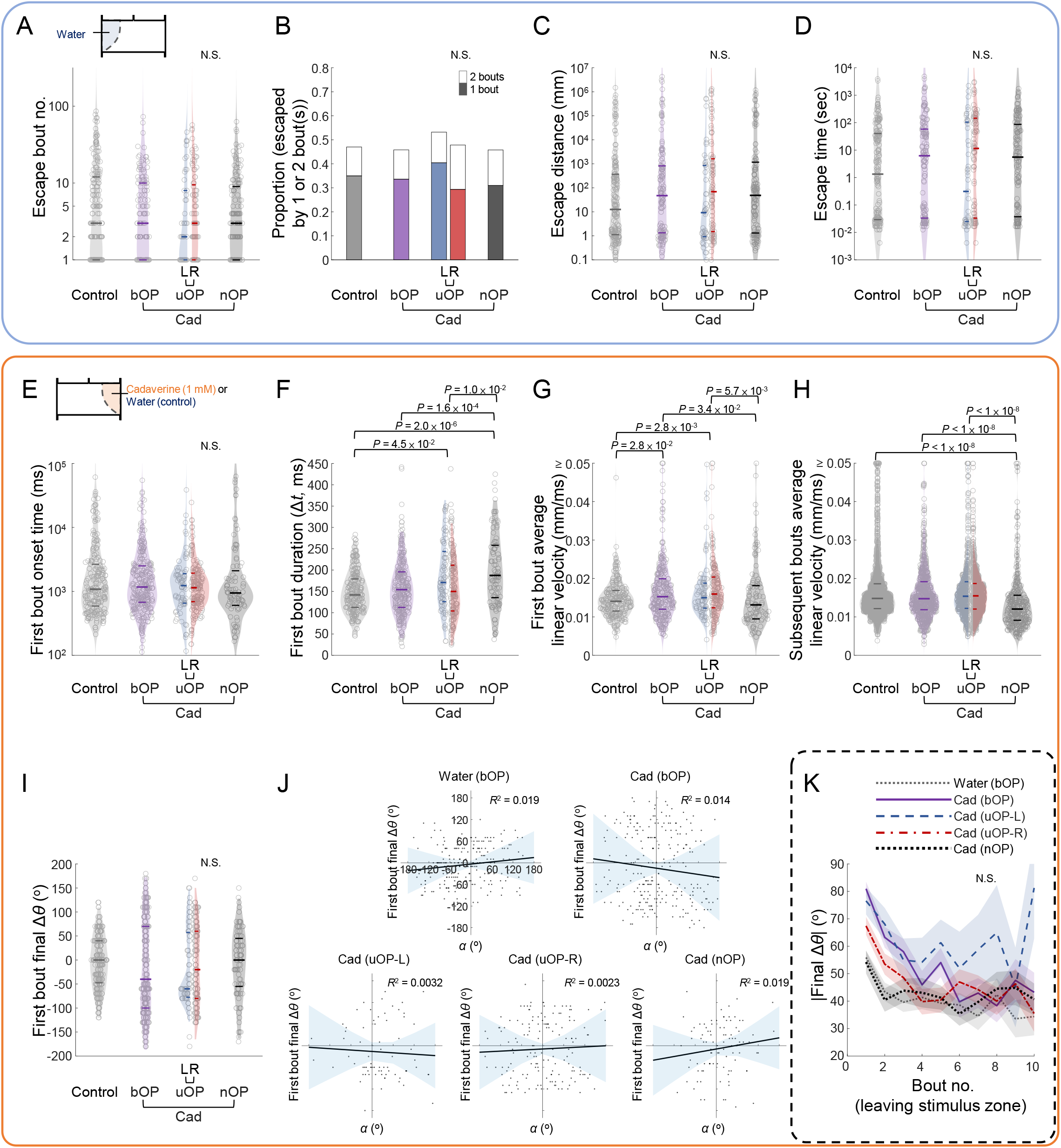
Performance metrics and kinemetric parameters in mirror zone and stimulus zone. **(A)** Bout number, **(B)** proportion of entry-to-exit events with only 1 or 2 bouts (N.S.: Non-significant with Chi-squared test comparing 2-bout event proportions), **(C)** distance travelled, and **(D)** time taken to escape the mirror zone in control assays (bOP larvae in water-only arena) and avoidance behavioral assays (bOP larvae, left OP-intact (L) or right OP-intact (R) uOP larvae, or nOP larvae in arenas with cadaverine stream in the stimulus zone). **(E) – (J)** Kinematic parameters of the first and subsequent bouts upon cadaverine encounter (as depicted in **Figure 2D** for first bouts), including **(E)** first bout onset time (relative to the end of entry bout), **(F)** first bout duration, **(G)** first bout average linear velocity, **(H)** subsequent bouts average linear velocity, and **(I)** final Δ*θ* for the first bouts. In **(A)**, **(C) – (E)**: Note that the parameters are plotted in log scales. In **(A) and (C) – (I)**: *P*-values: Kruskal–Wallis test with Tukey’s post-hoc test. N.S.: Non-significant. Horizontal lines indicate the medians, 75 and 25 percentiles for each group. **(J)** First turn final Δ*θ* plotted against incidence angle (*α*) on zone entry (also see **Figure 2D**). **(K)** |Final Δ*θ*| vs. bout number after leaving cadaverine zone (line: mean; shadow: SEM). | | denotes absolute value. N.S.: Non-significant with Mann-Kendall trend test.

**Supplementary Figure 3.**
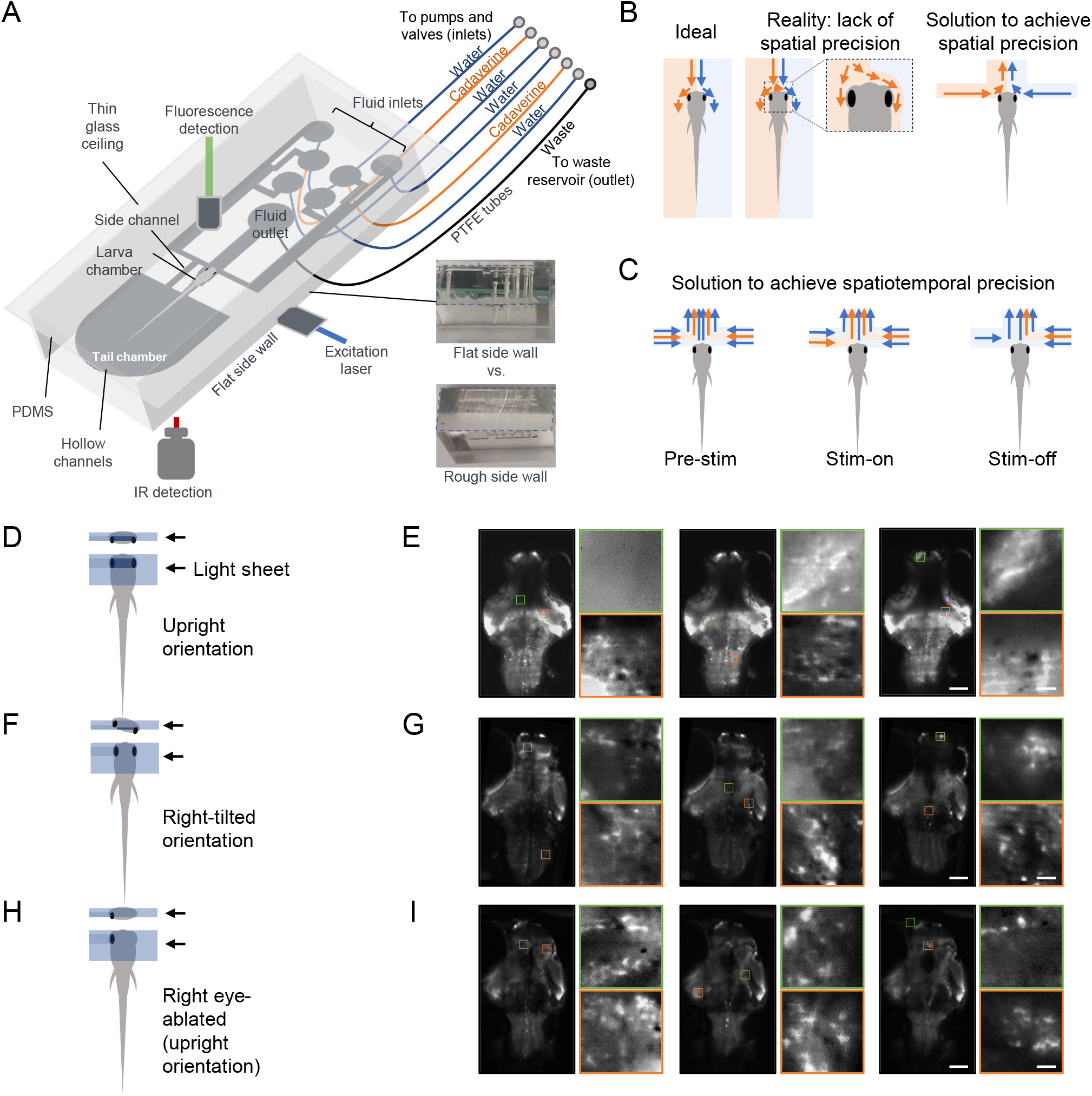
Detailed schematics, principles and imaging configurations for spatiotemporally precise nasal stimulation and whole-brain imaging. **(A)** Detailed schematics of the PDMS-based microfluidic device, that is compatible with (i) the perpendicular orientations of the excitation and detection optics of the custom-built light sheet microscope for whole-brain calcium imaging, and (ii) IR imaging module for the monitoring of stimulus-associated tail flicking behavior. The thin glass ceiling and the flat side wall (see blue dashed lines in the photos that outline the side surfaces for comparison, also see **Methods**) ensure undisturbed light propagation along the detection and excitation arms, respectively. Fluids are carried into and away from the microfluidic device via PTFE tubes, with inlets connected to valves and syringe pumps, and outlet to a waste reservoir. A long working distance detection objective (Nikon Plan Fluorite, ×10, N.A. 0.3, 16 mm WD) was used to accommodate the setup. **(B)** Left panel: Schematic depiction of the ideal segregation of different fluidic streams delivered from the front for unilateral OP stimulation with spatial precision. Light blue: water stream. Light orange: chemical stream. Arrows indicate the flow directions of the fluidic streams. Middle panel: In practical tests, we encountered unavoidable mixing of the fluidic streams flowing parallel from the front towards the larval zebrafish subject. This caused undesired spillover of chemical stimulus to the contralateral OP. Right panel: The problem was solved by directing fluidic streams from the sides to the OPs. The opposite directions of the streams that converge at the midline provided the necessary opposing forces to prevent spillover and achieve the required spatial precision of chemical stimulation. **(C)** The layout of fluidic streams that was eventually implemented to permit spatiotemporally precise chemical stimulation of unilateral OP. Left panel: At baseline (Pre-stim), the laminar water streams insulate the OPs from the chemical streams. Middle panel: By valve closure, the insulating water stream is removed (left side illustrated), and the chemical stream reaches the unilateral OP with a rapid rise time (Stim-on). Right panel: When the chemical stream is removed by valve closure, only the remaining water stream reaches the OP (Stim-off). Bilateral stimulation is achieved by undergoing the same sequence of fluid stream changes on both sides. **(D)** With an upright orientation, the right eye occludes the light sheet and results in poor illumination of the brain regions between the eyes. Upper panel: view from the tail of the larva. Lower panel: top view. Blue shadow represents the incoming light sheet with the direction indicated by the black arrows (i.e., from right to left). Areas with poor excitation are highlighted in darker colors. **(E)** Example images acquired in upright orientation from different planes. For each image set, the two images on the right are enlarged and contrast-adjusted images from the marked areas of the corresponding large field-of-view (FOV) images. Scale bars (white): 100 μm for the large FOV images and 10 μm for the zoomed-in images. **(F)** and **(G)**, similar to **(D)** and **(E)** but imaging with tilting by up to 20°to the right to circumvent the occlusion of light sheet entry by the right eye, thereby avoiding or minimizing poorly imaged regions. **(H)** and **(I)**, similar to **(D)** and **(E)** but imaging after surgical ablation of the right eye (12 – 24 hours prior to imaging).

**Supplementary Figure 4.**
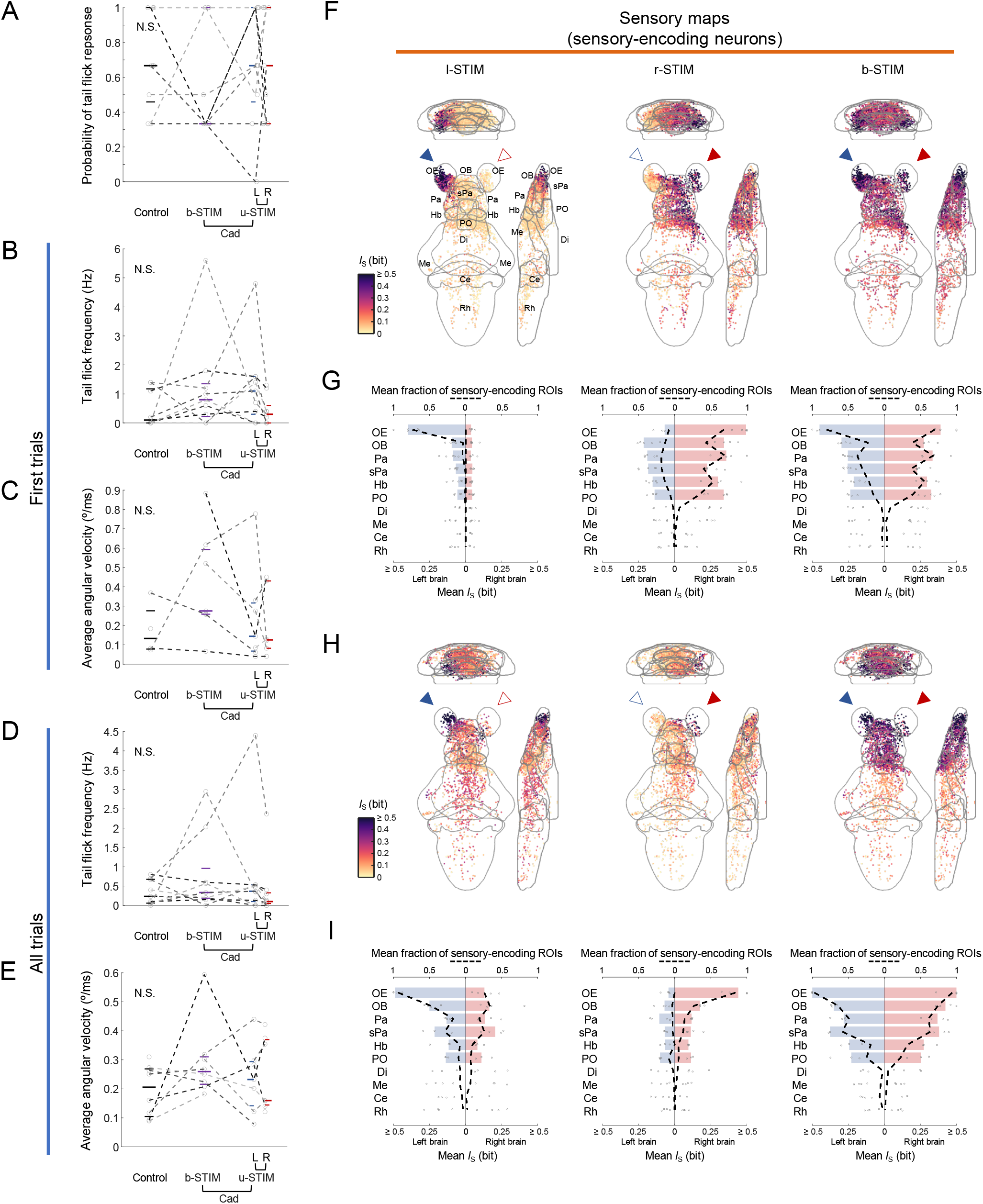
Motor responses and brainwide activation maps of larval zebrafish during spatiotemporally precise cadaverine stimulation. **(A)** Probability of tail flick response in the imaging device for each stimulus condition. **(B)** Tail flick frequency and **(C)** first-bout average angular velocity in the first trials of each stimulus condition. In all plots, one dot represents the mean of a given parameter of a larval subject in each stimulus condition. Data points from the same larval subjects are connected by dashed lines. Horizontal lines indicate the medians, 75 and 25 percentiles for each condition. **(D)** and **(E)**, similar to **(B)** and **(C)** respectively but with all bouts of all trials of each stimulus condition included. In **(A)** – **(E)**, N.S.: not significant with Kruskal–Wallis test, *n* = 9 behaviorally responsive larvae. **(F)** Mutual information (*I*_S_) maps pooled across 5 right eye-ablated larvae with biased responses (see **Methods**). Mean intensity projections of *I*_S_ between the calcium signals of sensory-encoding ROIs and cadaverine stimulus profile of l-STIM (left panel), r-STIM (middle panel) or b-STIM (right panel) are shown. Solid triangles mark the corresponding OP(s) stimulated. Abbreviations of brain regions: same as in **Figure 4**. **(G)** The distributions of mean *I*_S_ of sensory-encoding ROIs in the different brain regions during l-STIM (left panel), r-STIM (middle panel) or b-STIM (right panel) among the larvae (*n* = 5). Regions with top six mean fraction of sensory-encoding ROIs with b-STIM are OE, OB, Pa, sPa, Hb and PO. Bars representing the medians of the mean *I*_S_ of sensory-encoding ROIs in these regions. Dashed lines indicate fractions of sensory-encoding ROIs in the different regions among the larvae. **(H)** and **(I),** similar to **(F)** and **(G)** but with data pooled across larvae with generally unbiased responses to l-STIM or r-STIM (*n* = 4, consisting of 3 right-tilted and 1 right eye-ablated larvae, see **Methods**).

**Supplementary Figure 5.**
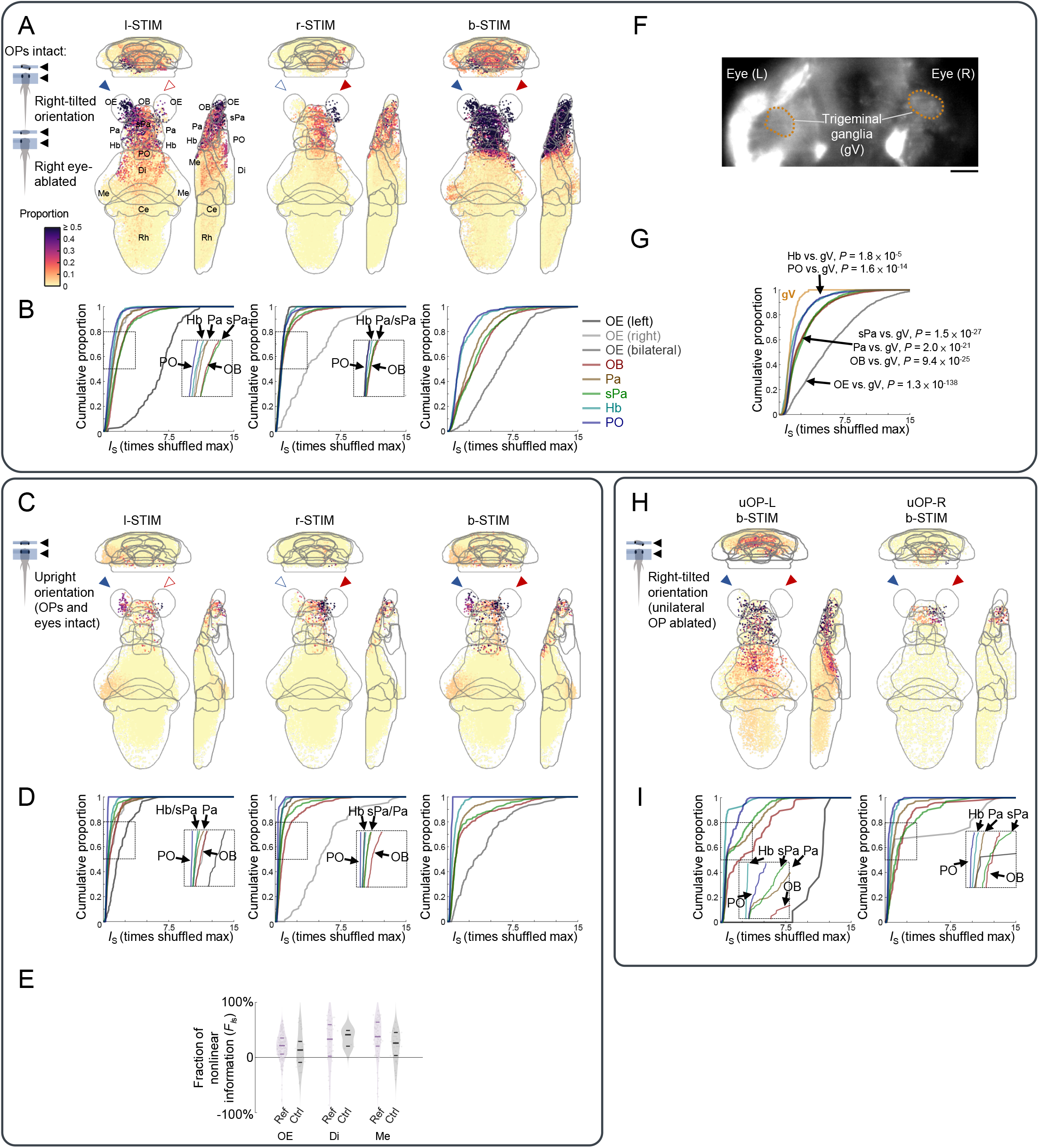
Brainwide regional activation analysis in upright orientation-imaged and unilateral OP-ablated larvae control groups, and comparison of rostral brain regions vs. trigeminal ganglia (gV) sensory encoding in cadaverine sensing. **(A)** Visualization of regional activation patterns during l-STIM (left panel), r-STIM (middle panel) or b-STIM (right panel) in the same larvae group shown in **Figure 5**, by color-coding all ROIs of each brain region by the corresponding regional proportions of sensory-encoding ROIs. Abbreviations of brain regions: same as in **Figure 4**. **(B)** Cumulative distributions of mutual information between the calcium signals of each ROI and cadaverine stimulus profile (*I*_S_ of l-STIM, r-STIM and b-STIM conditions, calculated as the number of times the maximum of the corresponding shuffled values) in the rostral brain regions. Insets correspond to the regions outlined by the dashed boxes. **(C)** and **(D)**: Similar to **(A)** and **(B)** but for a control larvae group (*n* = 4) with intact OPs and eyes imaged in an upright configuration. **(E)** Distributions of the fraction of nonlinear information (*F*_*I*s_) of ROIs in OE, Di and Me, in the larvae group shown in **(A)** (Ref) and the upright orientation-imaged larvae shown in **(C)** (Ctrl). Horizontal lines indicate the medians, 75 and 25 percentiles for each brain region. *P*-values: Wilcoxon rank-sum test. N.S.: Non-significant. **(F)** Example image plane showing the trigeminal ganglia (gV, outlined in orange dotted lines) in a larval zebrafish. Scale bar: 50 μm. **(G)** Cumulative distributions of mutual information between the calcium signals of each ROI and cadaverine stimulus profile (maximum *I*_S_ of l-STIM, r-STIM and b-STIM conditions) in gV (orange) and six rostral brain regions, in the larvae group shown in **(A)**. *P*-values: Two-sample Kolmogorov-Smirnov test comparing gV vs. each of the brain regions. **(H)** Visualization of regional activation patterns during b-STIM in left OP-intact (left panel, uOP-L, *n* = 2) and right OP-intact (right panel, uOP-R, *n* = 2) larvae, by color-coding all detected ROIs of each brain region by the corresponding regional proportions of sensory-encoding ROIs. **(I)** Cumulative distributions of mutual information between the calcium signals of each ROI and cadaverine stimulus profile (*I*_S_ of b-STIM condition) in six rostral brain regions of the same larvae group as **(H)**.

**Supplementary Figure 6.**
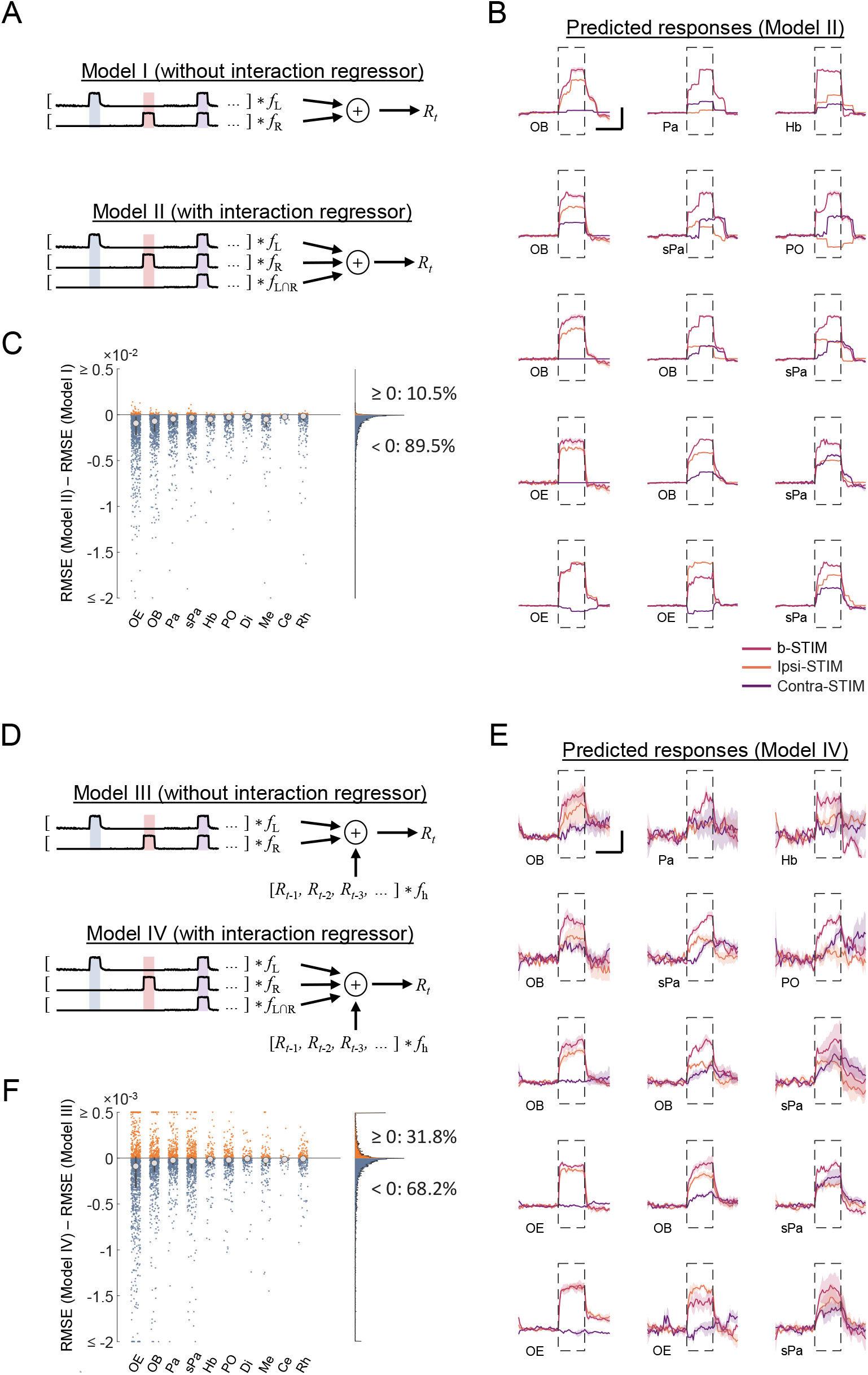
Generalized linear model (GLM) fitting to neuronal responses. **(A)** Two GLMs using only stimulus regressors for the individual OPs (Model I) or with the addition of a regressor to account for input interactions (Model II). *f*_L_, *f*_R_ and *f*_L∩R_ represent kernels for convolving with the l-STIM, r-STIM stimulus profiles and interaction regressors fitted to the models, respectively. Colored shadows indicate the chemical stimulus delivery windows. **(B)** Example trial-averaged predicted responses to ipsilateral (ipsi-STIM, orange), contralateral (contra-STIM, violet) or bilateral (b-STIM, cherry) OP stimulation of the same set of ROIs shown in **Figure 5D** with Model II (i.e., with interaction regressor). Shadow shows the SEM for each trace. **(C)** The distribution of root-mean-square error (RMSE) difference between Model II and Model I for individual ROIs in the different brain regions (*n* = 9 larvae). Large dots, upper and lower limits of lines: medians, 75 and 25 percentiles, respectively. Right panel: The distribution of RMSE difference pooled across regions. The proportions of ROI with RMSE(Model II) – RMSE(Model I) ≥ 0 (favoring Model I) and RMSE(Model II) – RMSE(Model I) < 0 (favoring Model II) are indicated. **(D), (E) and (F)**: Similar to **(A) – (C)**, but Model III and Model IV both have an additional regressor corresponding to the activity history of each ROI to be convolved with its fitted activity history kernel (*f*_h_). Example trialaveraged predicted responses of the same set of ROIs shown in **Figure 5D** with Model IV are shown in **(E)**. Dashed rectangle in **(B)** and **(E)** indicates stimulus window. Scale bars in **(B)** and **(E)**: 10 seconds (horizontal) and 0.5 normalized *dF*/*F* (vertical).

**Supplementary Figure 7.**
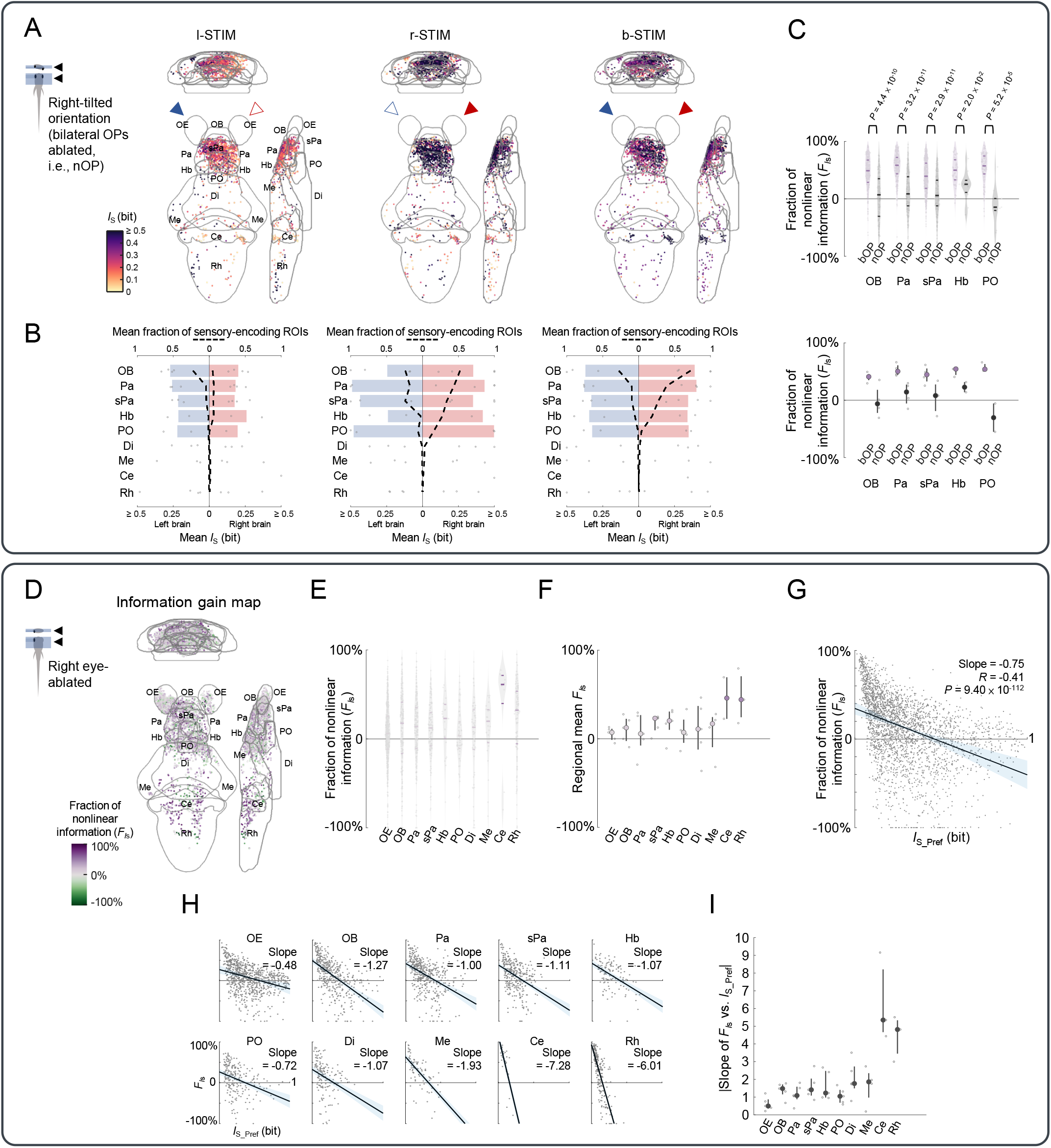
Brainwide sensory-encoding maps in bilateral OP-ablated larvae, and bilateral nasal input integration properties in the subset of right eye-ablated larvae with stronger response to r-STIM than l-STIM. **(A)** Mean intensity projections of *I_S_* between the calcium signals of sensory-encoding regions-of-interest (ROIs) and cadaverine stimulus profile of l-STIM (left panel), r-STIM (middle panel) or b-STIM (right panel) in larvae with bilateral OPs ablated (i.e., null OP-intact or nOP larvae, *n* = 3). Solid triangles mark the side of cadaverine stimulation. Abbreviations of brain regions: same as in **Figure 4**. **(B)** The distributions of mean *I*_S_ of sensory-encoding ROIs in the different brain regions during l-STIM (left panel), r-STIM (middle panel) or b-STIM (right panel) in the nOP larvae. Bars representing the medians of the mean *I*_S_ of sensory-encoding ROIs in the respective regions. Dashed lines indicate fractions of sensory-encoding ROIs in the different regions among the larvae. **(C)** Upper panel: Distributions of the fraction of nonlinear information (*F*_*I*s_) of ROIs in OB, Pa, sPa, Hb and PO, in bOP larvae (i.e., larvae with intact OPs, same dataset as in **Figure 5**) and nOP larvae groups. Horizontal lines: medians, 75 and 25 percentiles. Lower panel: Regional means of *F*_*I*s_, with each small dot representing the value from one larval zebrafish. Large dots, upper and lower limits of lines: medians, 75 and 25 percentiles, respectively. *P*-values: Wilcoxon rank-sum test. **(D)** Mean intensity projection maps of fraction of nonlinear information (*F*_*I*s_) in right eye-ablated larvae (*n* = 5). **(E)** Distributions of the *F*_*I*s_ of individual ROIs in the different brain regions. Horizontal lines: medians, 75 and 25 percentiles. **(F)** Regional means of *F*_*I*s_, with each small dot representing the value from one larval zebrafish. Large dots, upper and lower limits of lines: medians, 75 and 25 percentiles, respectively. **(G)***F*_*I*s_ versus *I*_S_ during preferred unilateral OP stimulation (*I*_S_Pref_, defined as *max*(*I*_S_l-STIM_, *I*_S_r-STIM_), where *I*_S_l-STIM_ and *I*_S_r-STIM_ denote *I*_S_ of l-STIM and r-STIM conditions, respectively) for ROIs pooled across regions and larvae. Black line and light blue shadow show line of best fit on linear regression with 95% confidence interval. Slope of the line of best fit, correlation coefficient (*R*) and *P*-value are shown. **(H)** Similar to **(G)** but with separate plot and linear regression performed for the ROIs of each brain region. Slopes of the lines of best fit are shown. **(I)** The absolute values of the slope of *F*_*I*s_ versus *I*_S_Pref_ for each brain region (i.e., |*F*_*I*s_ vs. *I*_S_Pref_|), with each small dot representing the value from one larval zebrafish. Large dots, upper and lower limits of lines: medians, 75 and 25 percentiles, respectively.

## Notes

### Competing Interest Statement

The authors have declared no competing interest.

### Summary of Updates

We added results from more control experiments and refined / updated the analysis.

